# Combined MEK and STAT3 inhibition reprograms stromal inflammation to overcome immunotherapy resistance in pancreatic cancer

**DOI:** 10.1101/2021.03.07.434236

**Authors:** Jashodeep Datta, Xizi Dai, Anna Bianchi, Iago De Castro Silva, Siddharth Mehra, Vanessa Garrido, Purushottam Lamichhane, Samara Singh, Zhiqun Zhou, Austin R. Dosch, Fanuel Messaggio, Yuguang Ban, Oliver Umland, Peter J. Hosein, Nagaraj S. Nagathihalli, Nipun B. Merchant

## Abstract

Pancreatic ductal adenocarcinoma (PDAC) is characterized by immune exclusion, pro-inflammatory polarization of cancer-associated fibroblasts (CAF), and resistance to immune checkpoint inhibition (ICI). We have previously demonstrated that reciprocally activated RAS/MEK/ERK and JAK/STAT3 pathways mediate therapeutic resistance, while combined MEK and STAT3 inhibition (MEKi+STAT3i) overcomes such resistance in preclinical models. We now show that combined MEKi+STAT3i not only alters stromal architecture but also uncovers stromal plasticity by revealing a substantial attenuation of *Il6/Cxcl1*-expressing secretory and *Lrrc15*-expressing myofibroblastic CAF phenotypes with a concomitant enrichment of *Ly6a/Cd34*-expressing CAF phenotypes exhibiting mesenchymal progenitor-like properties via single-cell RNA sequencing in *Ptf1a^cre/+^;LSL-Kras^G12D/+;^Tgfbr2^flox/flox^*(PKT) mice. This remodeling of CAF heterogeneity is associated with reprogramming of immunosuppressive myeloid populations and enhanced trafficking of CD8^+^ T-cells which exhibit a distinct effector transcriptional program. These MEKi+STAT3i-mediated repercussions are in part CAF-dependent, since CRISPR/Cas9 genetic silencing of CAF-restricted *Mek1*/*Stat3* mitigates inflammatory CAF polarization and myeloid infiltration *in vivo*. Addition of MEKi+STAT3i to PD-1 blockade overcomes ICI resistance by significantly augmenting anti-tumor responses and dramatically improving survival in PKT mice compared with anti-PD-1 monotherapy. The addition of MEKi+STAT3i to PD-1 blockade not only augments the recruitment of activated and memory T-cell populations, but also improves their degranulating capacity and functional cytotoxicity compared to PD-1 blockade alone. Importantly, treatment of a patient with chemotherapy-refractory metastatic PDAC with MEKi (Trametinib), STAT3i (Ruxolitinib), and PD-1 inhibitor (Nivolumab) was well-tolerated and yielded clinical benefit. These data uncover a novel paradigm in which combined MEKi+STAT3i reprograms stromal inflammation and immune tolerance to overcome immunotherapy resistance in PDAC.

## INTRODUCTION

Pancreatic ductal adenocarcinoma (PDAC) remains a major therapeutic challenge due to its intrinsic and acquired chemoresistance. The major contributors to therapeutic resistance that have been difficult to overcome in PDAC are mutations in the *KRAS* oncogene (*1*), the presence of a dense desmoplastic stroma that acts as a barrier to drug delivery and effector immune cell infiltration (*2*), and the immunosuppressive tumor microenvironment (TME) that renders the tumor resistant to immune checkpoint inhibition (ICI) immunotherapy (*3*).

Despite unprecedented success in other solid tumors (*4*), clinical trials utilizing single or dual ICI have been decidedly unsuccessful in PDAC patients (*5, 6*). The failure of ICI in PDAC has been attributed not only to the fibroinflammatory desmoplastic stroma in PDAC that promotes T-cell exclusion, but also to the establishment of an immunosuppressive myeloid ecosystem in the TME. Cellular elements of this ecosystem comprise tumor-associated macrophages (TAM) and myeloid-derived suppressor cells (MDSC), which induce T-cell suppression through diverse mechanisms (*7*). A critical unmet need in the field of PDAC immunotherapy, therefore, is the discovery of novel therapeutic strategies to reprogram its tolerogenic immune microenvironment.

Beyond stromal desmoplasia and the biomechanical properties of extracellular matrix (ECM) components that impede drug delivery and T-cell infiltration (*8*), emerging evidence implicates functional heterogeneity in the cancer-associated fibroblast (CAF) compartment as a key stromal-mediated mechanism of immunotherapy resistance in PDAC (*9, 10*). In particular, IL1-mediated polarization of CAFs toward a secretory phenotype, characterized by elaboration of IL-6, CXCL1, CXCL12, and LIF (*11, 12*), propagates pro-inflammatory tumor-stromal-immune crosstalk in the PDAC TME that drives therapeutic resistance (*13*). These tumor-permissive inflammatory CAFs (iCAF)—which co-exist in a dynamic equilibrium with other functionally divergent CAF sub-populations such as myofibroblastic CAFs (myCAFs) and antigen-presenting CAFs (apCAFs) (*14, 15*)—beckon immunosuppressive TAMs and MDSCs to the TME, dampen anti-tumor adaptive immunity, and promote immune evasion in PDAC (*11, 13, 16, 17*). These underlying complexities in CAF immunobiology, coupled with the clinical failure of *broad-based* stromal-depleting strategies in PDAC patients (*18, 19*), warrant the development of more nuanced therapies to mitigate tumor-permissive inflammatory CAF programming while preserving its tumor-restraining counterparts.

*KRAS* mutations—another key contributor to therapeutic resistance—are not only the predominant oncogenic driver in over 90% of PDAC, but also promote an inflammatory program that establishes immune privilege in the TME (*1, 20*). Since targeting *KRAS* has remained elusive, our approach has been to target downstream effectors of RAS through MEK inhibition (MEKi). However, clinical trials of MEKi have been unsuccessful in PDAC (*21*), likely due to the activation of resistance mechanisms. We have shown that MEK inhibition (MEKi) results in reciprocal activation of STAT3 signaling, which confers therapeutic resistance and continued PDAC cell growth. Combined JAK/STAT3 inhibition (STAT3i) and MEKi (MEKi+STAT3i) overcomes this resistance mediated by parallel feedback loop activation, attenuates tumor growth in patient-derived xenograft models, and improves survival in the aggressive *Ptf1a^cre/+^;LSL-Kras^G12D/+;^Tgfbr2^flox/flox^*(PKT) genetically engineered mouse model (GEMM) (*22, 23*).

In the present study, we show that combined MEKi+STAT3i uncovers stromal plasticity by attenuating tumor-resident CAFs from *Il6/Cxcl1*-expressing secretory phenotypes and enriching for *Ly6a/Cd34*-expressing CAF phenotypes with mesenchymal progenitor-like properties via single-cell RNA sequencing (scRNAseq) in PKT mice. This remodeling of CAF heterogeneity following MEKi+STAT3i treatment is associated with a striking attenuation in and reprogramming of TAMs, as well as enhanced trafficking of CD8^+^ T-cells which exhibit a distinct effector and anti-apoptotic transcriptional program. These immune repercussions are in part CAF-dependent, since CRISPR/Cas9 genetic silencing of CAF-restricted *Mek1/Stat3* results in attenuation of iCAF polarization and MDSC/TAM infiltration *in vivo.* Importantly, the addition of MEKi+STAT3i to PD-1 blockade overcomes ICI resistance by significantly augmenting anti-tumor responses and dramatically improving survival in PKT mice, compared with vehicle treatment or anti-PD-1 monotherapy. The addition of MEKi+STAT3i to PD-1 blockade not only augments the recruitment of activated and memory T-cell populations, but also improves their degranulating capacity and functional cytotoxicity. Finally, we demonstrate the clinical efficacy and tolerability of combined Trametinib (MEKi), Ruxolitinib (STAT3i), and Nivolumab (anti-PD1) treatment in a patient with chemotherapy-refractory PDAC, providing evidence of its translatability in PDAC patients.

## RESULTS

### Combined MEK and STAT3 inhibition remodels stromal fibrosis and attenuates inflammatory CAF phenotypes in a fibroblast-dependent manner

We have previously shown that combined MEKi+STAT3i decreases tumor burden and improves survival in PKT mice (*22*). We extend these results here to show that combined MEKi+STAT3i significantly remodels the tumor stroma as evidenced by decreased trichrome blue, smooth muscle actin (α-SMA), and Sirius Red staining of pancreatic tumor sections in PKT mice (**Fig. 1A**). Moreover, MEKi+STAT3i significantly reduces intratumoral mucin content (Alcian Blue staining) and cellular proliferation (Ki67), as well as increases microvessel density (CD31 staining) compared with vehicle treatment (**Fig. S1**).

**Figure 1.**
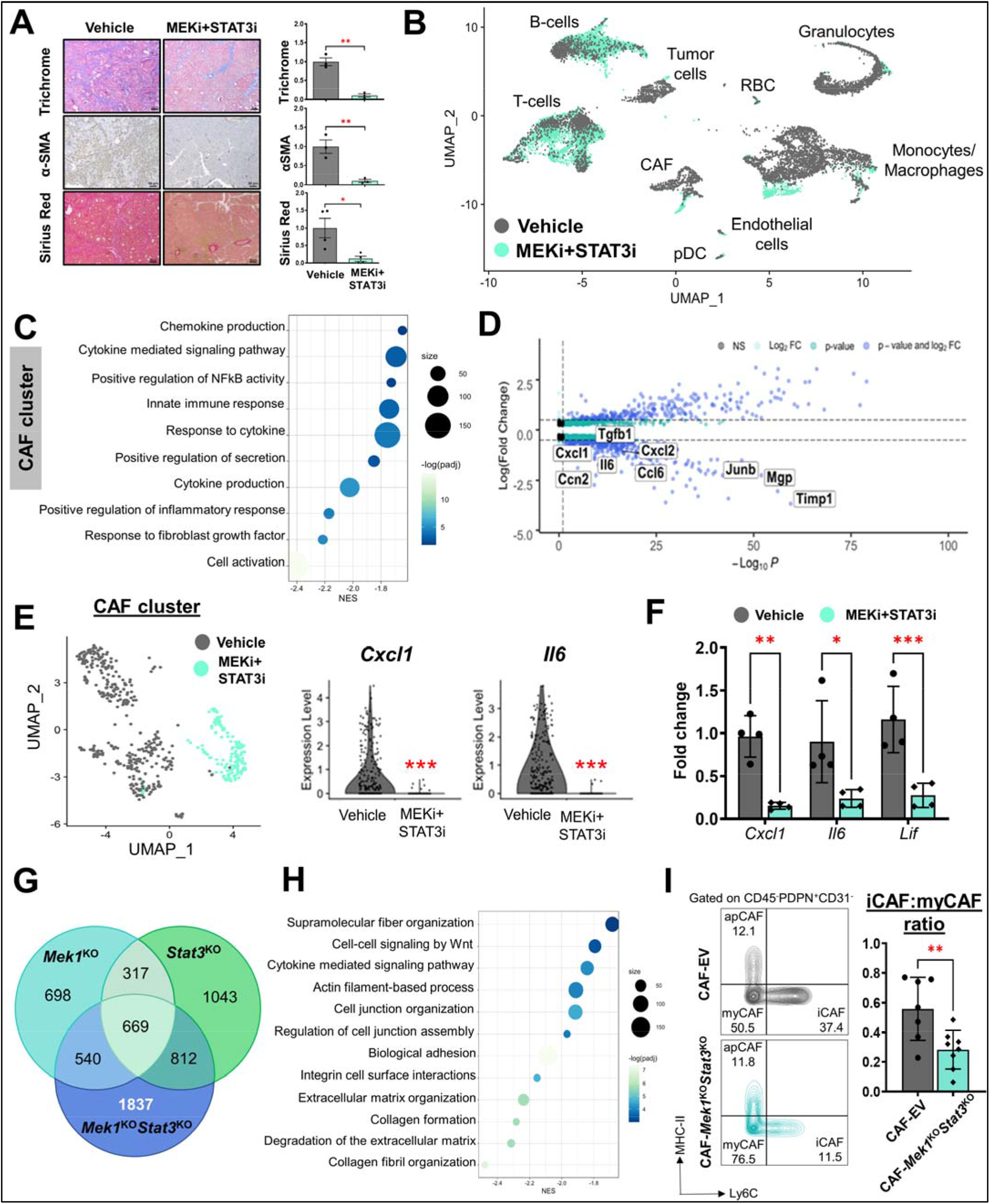
Combined MEK and STAT3 inhibition remodels stromal fibrosis and attenuates inflammatory fibroblast phenotypes in the TME in a CAF-dependent manner. (**A**) Trichrome blue, α-SMA, and Sirius Red staining in tumor sections from PKT mice treated with vehicle or MEKi+STAT3i for 4 weeks; relative areas of positive staining of respective markers from tissue sections in vehicle- and MEKi+STAT3i-treated mice are indicated in adjacent histograms (scale bar=50 µm). Data are shown as mean ± SEM; (**B**) Concatenated UMAP plot showing annotated clusters from 12,680 single cells undergoing RNA sequencing from vehicle- and MEKi+STAT3i-treated PKT mice (n=3 each); (**C**) Bubble plot representing pathway enrichment analysis performed on genes differentially downregulated in MEKi-STAT3i treated CAF cluster using fgsea (log fold change (FC)>1). Reactome, KEGG, and GO pathways with *p-* adjusted value<0.05 are displayed with normalized enrichment score (NES) indicated on x-axis; (**D**) Volcano plot of select differentially regulated genes related to innate immune response and stromal organization in MEKi+STAT3i-treated vs. vehicle-treated CAF single-cell transcriptomes. Vertical dashed line indicates an adjusted *p-*value=0.1, horizontal dashed lines indicate an absolute log_2_FC=0.5. Transcripts achieving non-significance (NS), p-value, log_2_FC, or p-value and log_2_FC significance threshold are indicated in adjoining legend; (**E**) CAF clusters in vehicle and MEKi+STAT3i-treated superimposed UMAP plot (*left*); violin plots depicting log expression level of *Cxcl1* and *Il6* genes in CAF single-cell transcriptomes comparing MEKi+STAT3i vs. vehicle-treated CAFs; (**F**) RT-qPCR analysis from whole tumor-derived RNA in MEKi+STAT3i-treated vs. vehicle-treated primary PKT tumors (n=4 mice/group) depicting relative fold change of *Cxcl1, Il6,* and *Lif* gene expression; (**G**) Venn diagram of differentially expressed genes in bulk RNA sequencing derived from KPC CAF cell lines that underwent CRISPR-Cas9 genetic silencing of either *Mek1*^KO^ alone, *Stat3*^KO^ alone or combined *Mek1*^KO^/*Stat3*^KO^. Numbers in each circle (intersection) represent the unique number of differentially regulated genes in respective comparisons; 1837 genes were differentially expressed in CAF-*Mek1*^KO^*Stat3*^KO^, and were utilized for further analysis; (**H**) Bubble plot representing pathway enrichment analysis performed on genes differentially downregulated in CAF-*Mek1*^KO^*Stat3*^KO^ cells. Reactome, KEGG, and GO pathways with p-adjusted value<0.05 are displayed with normalized enrichment score (NES) indicated on x-axis; (**I**) Representative contour plots of CD45^-^PDPN^+^CD31^-^ cells showing MHC-II and Ly6C gates to indicate iCAF (Ly6C^+^MHC-II^-^), myCAF (Ly6C^-^MHC-II^-^), and apCAF (Ly6C^+^MHC-II^+^) populations from experiments in which tumors were isolated from orthotopically injected C57BL/6 mice co-injected with KPC6694c2 tumor cells and either CAF-EV or CAF-*Mek1*^KO^/*Stat3*^KO^ CAF (ratio 1:9 respectively) after 2 weeks (n=7-8 mice/group). Adjacent histogram shows quantification of iCAF:myCAF ratio at endpoint analysis. ns, not significant; *, p<0.05; **, p<0.01; ***, p<0.001.

Based on these striking findings, we hypothesized that MEKi+STAT3i treatment may promote stromal remodeling and exert its antitumor effects by unmasking CAF plasticity and functionally reprogramming the CAF compartment. To examine mechanistic underpinnings of MEKi+STAT3i-induced stromal remodeling *in vivo*, scRNAseq was performed in PKT mice—a PDAC GEMM characterized by dense stromatogenic response and immune exclusion (*23, 24*)—treated with MEKi+STAT3i or vehicle. Using differentially expressed gene signatures, we attributed clusters to their putative identities (**Fig. S2A**). Specifically, CAF clusters were nominated by exclusive expression of *Col1a1, Col1a2, Pdpn,* and *Fap* (**Fig. S2B**) (*25, 26*). Uniform Manifold Approximation and Projection (UMAP), an unsupervised dimensionality reduction clustering algorithm, was used to display discrete intratumoral cellular compartments in PKT mice (**Fig. 1B**).

Compared with vehicle treatment, combined MEKi+STAT3i resulted in substantial reprogramming of both the CAF and tumor-cell transcriptomes (**Fig. S3A&B**). Deeper investigation of the differentially expressed single-cell transcriptomes in the CAF compartment of MEKi+STAT3i- vs. vehicle-treated mice revealed significant downregulation of signaling pathways related to inflammatory cytokine/chemokine response, innate immune trafficking, NF-κB signaling, and extracellular matrix (ECM) deposition (**Fig. 1C**). Moreover, several genes implicated in promoting a pro-inflammatory secretome and/or innate immune cell recruitment (*Cxcl2, Ccl6),* peripheral immune tolerance (*Tgfb1*), and stromal organization/equilibrium (*Mgp, Junb, Timp1*) were strongly differentially under-expressed in MEKi+STAT3i-treated, compared with vehicle-treated, CAF transcriptomes (**Fig. 1D**). In particular, expression of iCAF-defining genes *Cxcl1* and *Il6* (*14*) was significantly reduced in single-cell CAF transcriptomes following MEKi+STAT3i treatment (**Fig. 1E**). These findings were validated by significant diminution of *Cxcl1* (P=0.002)*, Il6* (P=0.01)*, and Lif* (P<0.001) expression in MEKi+STAT3i-treated vs. vehicle-treated PKT tumors via qPCR analysis in whole tumor-derived RNA (**Fig. 1F**).

Next, to determine if MEKi+STAT3i attenuates iCAF skewness in a fibroblast-dependent manner, we performed CRISPR/Cas9-enabled genetic editing of *Mek1* alone (*Mek1*^KO^), *Stat3* alone (*Stat3*^KO^), and dual *Mek1/Stat3* (*Mek1*^KO^*Stat3*^KO^) in FAP^+^PDPN^+^ *Kras^G12D/+^;Trp53^fl/+^;Pdx-^Cre^* (KPC) tumor-derived CAFs, and confirmed target silencing using western blotting (**Fig. S4A**). Characterizing these cells via bulk RNA sequencing revealed a distinct transcriptome in *Mek1*^KO^*Stat3*^KO^ CAFs comprising 1837 unique genes (**Fig. 1G**) that collectively reflected downregulation of pathways related predominantly to focal adhesion/ECM deposition, collagen organization, and cytokine signaling (**Fig. 1H**; **Fig. S4B**). Compared to empty vector control-CAFs (CAF-EV), the *in vitro* secretome CAF-*Mek1*^KO^*Stat3*^KO^ demonstrated reduction in several myeloid chemoattractants, such as IL-6, CXCL1, GM-CSF, and CCL2 (**Fig. S4C**), recapitulating the effect of MEKi+STAT3i on scRNAseq CAF transcriptomes in PKT mice.

Next, to determine the effect of CAF-specific *Mek1/Stat3* silencing on CAF skewness *in vivo,* orthotopic co-injection of KPC6694c2 tumor cells with either KPC CAF-EV or CAF-*Mek1*^KO^*Stat3*^KO^ (1:9 ratio) in syngeneic C57Bl/6 mice resulted in significant reduction of iCAF (CD45^-^CD31^-^EpCAM^-^PDPN^+^Ly6C^+^MHC-II^-^) to myCAF (CD45^-^CD31^-^EpCAM^-^PDPN^+^Ly6C^-^MHC-II^-^) cellular ratios by flow cytometry in CAF-*Mek1*^KO^*Stat3*^KO^ vs. CAF-EV tumors (P=0.009; **Fig. 1I**). Collectively, these data suggest that MEKi+STAT3i remodels stromal inflammation and attenuates iCAF phenotypes in a fibroblast-dependent manner *in vivo*.

### Combined MEK and STAT3 inhibition uncovers stromal plasticity and enriches for a CAF phenotype with mesenchymal progenitor-like properties

To illustrate how MEKi+STAT3i shapes CAF transcriptional heterogeneity in the PDAC TME, CAF-designated barcoded events on scRNAseq were projected by UMAP and revealed the presence of four discrete sub-clusters (**Fig. 2A**) harboring distinct gene expression profiles (**Fig. S5A**). While CAF2, CAF3, and CAF4 sub-populations dominated the CAF landscape in vehicle-treated mice, MEKi+STAT3i treatment revealed an enrichment in the CAF1 cluster while substantially attenuating CAF2, CAF3, and CAF4 populations (**Fig. 2B**). Given this striking MEKi+STAT3i-induced remodeling of the fibroblast microenvironment, we further investigated the transcriptional and functional divergence between CAF1 and CAF2/CAF3/CAF4 subsets. Principal component analysis based on the most highly variable genes, as measured by variance-stabilizing transformation (**Fig. S5B**), and matrix plots of population concordance, as measured by Pearson’s correlation and Euclidean distances prior to hierarchical clustering of differentially expressed genes between CAF1-4 sub-clusters (**Fig. 2C**), revealed the largest degree of transcriptional separation between CAF1 and CAF2-4.

**Figure 2.**
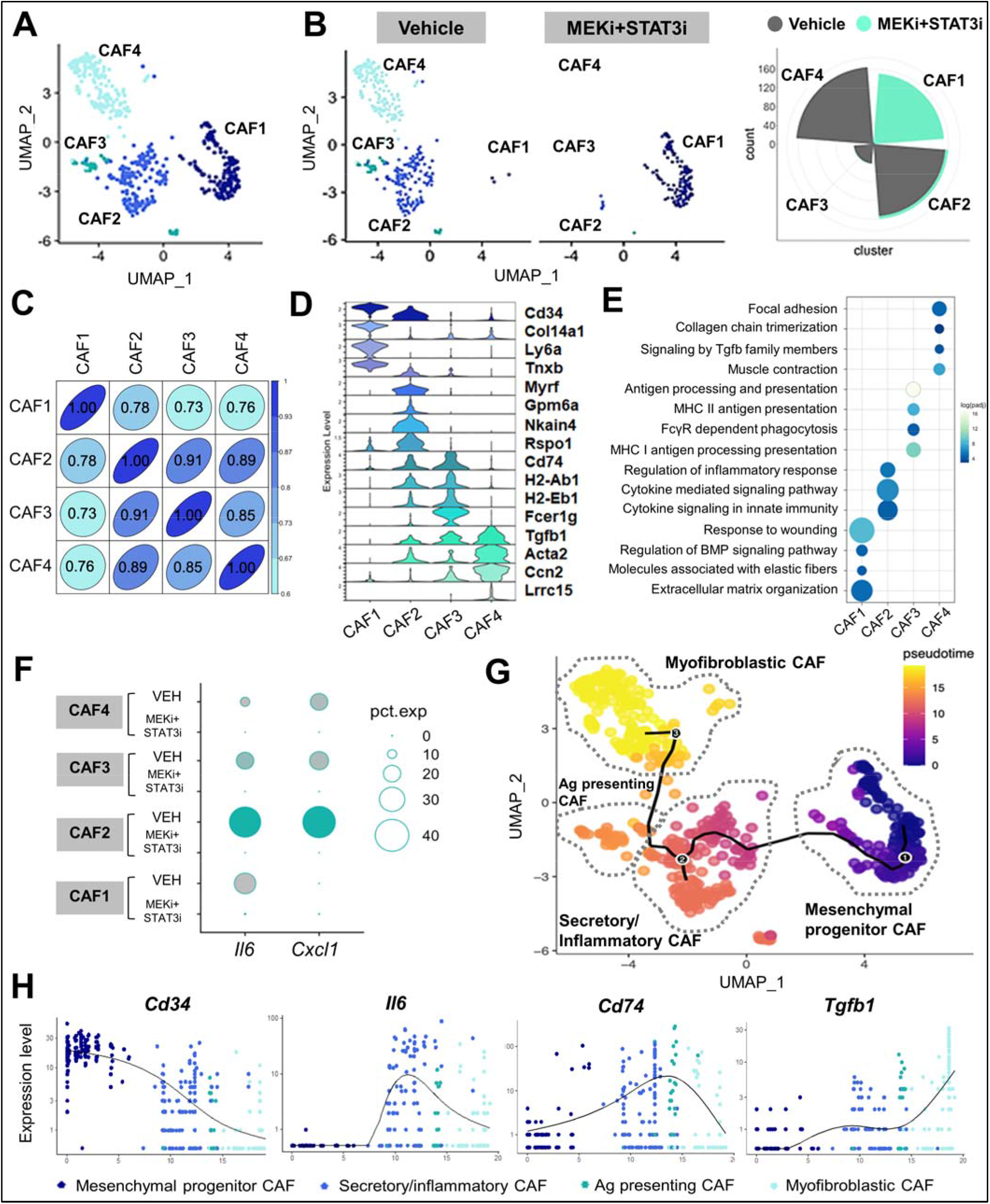
Combined MEK and STAT3 inhibition uncovers stromal plasticity and emergence of a CAF population with mesenchymal progenitor-like properties. **(A)** Superimposed UMAP plot highlighting emergent CAF sub-clusters in vehicle- and MEKi+STAT3i treated PKT tumors subjected to single-cell RNA sequencing (scRNAseq); **(B)** Adjacent UMAP plots and circumplex plots showing absolute cell count of each CAF sub-cluster in vehicle- and MEKi+STAT3i-treated mice, respectively; **(C)** Pearson correlation analysis of transcriptional divergence between CAF sub-clusters in the combined vehicle- and MEKi+STAT3i CAF Seurat object. 1.00=complete correlation, 0.00=no correlation; **(D)** Stacked violin plots depicting expression level of select genes uniquely expressed in each CAF sub-cluster CAF1-4; **(E)** Bubble plot of pathway enrichment analysis highlighting the strongly differentially enriched pathways in each CAF cluster using fgsea (log(FC)>1. Reactome, KEGG, and GO pathways with adjusted p-value<0.05 were selected. **(F)** Dot plot depicting the expression of iCAF-defining genes *Il6* and *Cxcl1* across different CAF sub-subsets comparing vehicle- and MEKi+STAT3i-treated cohorts; **(G)** Single-cell lineage trajectory analysis performed using Monocle3 pipeline colored by Pseudotime timestamps (adjoining legend) in CAF-only cluster depicting 4 major putative CAF cellular sub-clusters in superimposed UMAP plot. The major trajectory is depicted by the continuous solid starting at (1), and two minor bifurcations indicated by branchpoints (2) and (3). CAF sub-clusters are nominated by gene expression identity and putative pathway enrichment into mesenchymal progenitor (CAF1), secretory/inflammatory (CAF2), antigen-presenting (CAF3), and myofibroblastic (CAF4); **(H)** Expression of lineage state-defining markers (*Cd34, Il6, Cd74, Tgfb1*) on Pseudotime scale across the CAF sub-clusters.

Closer inspection of the differentially expressed single-cell transcriptomes in CAF1-4 sub-clusters revealed distinct transcriptional programs that reflected *functional* specialization. Differentially expressed genes (using log(fold-change)>0.5, p-adj<0.05 threshold) were subjected to pathway enrichment analysis to determine functional signatures defining each CAF subset. While CAF lineage-specific markers *Fap, Pdpn*, and *Col1a1* were expressed in all these sub-clusters (**Fig. S1B**), CAF1—the most transcriptionally disparate from the other three sub-clusters—was strongly enriched in the mesenchymal fibrocytic marker *Cd34*, suggesting its putative origin from a bone marrow-derived hematopoietic/mesenchymal progenitor-like lineage; *Ly6a* (or *Sca-1)*, a mesenchymal stromal/stem cell (MSC) marker detected early in pancreatic neoplasia but progressively lost during tumorigenesis (*26*); and *Col14a1*, a discriminatory marker for matrix fibroblasts (*27*) (**Fig. 2D**). This distinct gene signature correlated with a concentration of pathways reflecting fundamental processes related to fibroblast biology, such as matrix synthesis/organization, response to wounding, and elastic fiber formation (**Fig. 2E**). Intriguingly, CAF1 strongly expressed the bone marrow-derived MSC marker Meflin (or *lslr)*, which has been recently proposed as a marker of tumor-restraining CAFs in PDAC (**Fig. S5C**) (*28*). Based on these features, CAF1 was nominated as *mesenchymal progenitor CAF*.

The CAF2 sub-cluster was significantly enriched for pathways associated with pro-inflammatory and immunomodulatory function, including cytokine signaling involved in innate immune responses (**Fig. 2E**). The concentration of secretory markers *Il6, Cxcl1,* and *Il33* in the CAF2 subset was consistent with its inflammatory transcriptional program (**Fig. S5D**). The global reduction in CAF-specific *Il6/Cxcl1* expression following MEKi+STAT3i (**Fig. 1E**), therefore, was driven predominantly by attenuation of CAF2 subpopulations in the TME (**Fig. 2F**). CAF2 was nominated as *secretory/inflammatory CAF*.

The CAF3 sub-cluster was enriched in genes related to antigen-presenting function *Cd74, H2-ab1, H2-Eb1* (**Fig. 2D**) and MHC class II-related pathways (**Fig. 2E**); interestingly, *Fcer1g*—a classical macrophage marker—and FcγR-dependent phagocytosis pathways were also enriched in CAF3, suggestive of a shared lineage with myeloid-derived antigen-presenting cells. Thus, CAF3 was nominated as *antigen-presenting CAF.* The CAF4 sub-cluster was enriched in the transcripts *Tgfb1, Acta2*, and *Ccn2* (**Fig. 2D**), indicative of canonical myofibroblastic functions such as smooth muscle contraction and focal adhesion, as well as enrichment of TGF-β signaling (**Fig. 2E**). Notably, the myCAF marker *Lrrc15*—recently implicated as a master regulator of CAF-mediated resistance to immunotherapy and key driver of T-cell exclusion in PDAC (*9*)—was exclusively expressed in CAF4 (**Fig. 2D**), and its expression abrogated in MEKi+STAT3i-treated PKT tumors (**Fig. S5E**). CAF4 was therefore nominated as *myofibroblastic CAF*.

### Pseudotime lineage reconstruction reveals differential developmental trajectories of CAF subsets following MEKi+STAT3i

We next investigated if the transcriptionally disparate CAF subsets uncovered by MEKi+STAT3i treatment had convergent or divergent developmental trajectories. Ordering of cells in pseudotime lineage reconstruction analysis (*29*) arranged CAFs into a major trajectory commencing with mesenchymal progenitor CAFs—transcriptionally similar to Hosein *et al*. FB2, progressively lost during PDAC tumorigenesis (*26*)—with two minor bifurcations: secretory/inflammatory CAFs (i.e., CAF2) and myofibroblastic CAFs (i.e., CAF4) (**Fig. 2G**). CAFs expressing MSC markers *Cd34, Ly6a, and Ly6c1* and matrix fibroblast-specific transcript *Col14a1* preferentially distributed at the beginning of all paths (**Fig. 2H**; **Fig. S6**), while the lineage-determining genes *Il6*, *Cd74*, and *Tgfb1* increased in density and expression levels progressively in CAFs nominated transcriptionally as secretory/inflammatory, antigen-presenting, and myofibroblastic, respectively (**Fig. 2H**; **Fig. S6**).

Interestingly, the enrichment of canonical mesothelial markers *Myrf*, *Gpm6a*, and *Nkain4* in the predicted lineage paths of secretory/inflammatory as well as antigen-presenting CAFs (**Fig. S6**) suggest a co-evolution of these populations from a shared mesothelial ontology. Expression of these mesothelial markers dissipated in the lineage paths at the 2^nd^ trajectory bifurcation toward myofibroblastic CAFs, revealing instead an enrichment in contractile genes *Acta2* and *Ccn*. Taken together, these data suggest that MEKi+STAT3i treatment uncovers distinct CAF plasticity by enriching for an MSC-like mesenchymal progenitor CAF subset while attenuating mesothelial- and myofibroblast-derived CAF populations in the PDAC TME.

### MEKi+STAT3i reprograms the immunosuppressive myeloid microenvironment and facilitates intratumoral T-cell trafficking in part via a CAF-dependent manner

Next, given the mitigation of IL-6/CXCL1-producing iCAF subsets following MEKi+STAT3i treatment, we examined if this reprogramming of stromal inflammation results in remodeling of the innate immune microenvironment *in vivo* (*30*). Indeed, flow cytometric analysis of PKT tumors (n=8-10 mice/arm) revealed a significant decrease in CD11b^+^ myeloid cells and F4/80^+^ macrophages (**Fig. S7A**).

To comprehensively assess the remodeling of the innate immune microenvironment associated with MEKi+STAT3i at single-cell resolution in PKT mice, we performed multiparametric immune profiling of pooled pancreatic tumor samples utilizing time-of-flight mass cytometry (CyTOF). Compared with vehicle treatment, MEKi+STAT3i, resulted in broad-based changes in the innate (CD11b^+^), adaptive (CD4^+^ and CD8^+^), and humoral (CD19^+^) immune compartments (**Fig. 3A**). In particular, compared with vehicle treatment, MEKi+STAT3i treatment resulted in a dramatic decrease in CD11b^+^ myeloid cells, F4/80^+^ macrophages, M2-like macrophages (F4/80^+^CD206^+^), and MDSCs (CD11b^+^F4/80^-^Ly6G^+^/Ly6C^+^) (**Fig. 3B**). Furthermore, scRNAseq of PKT tumors analysis confirmed these broad-based changes in immune subpopulations following MEKi+STAT3i compared with vehicle treatment (**Fig. 3C**). Beyond decreased abundance of monocyte/macrophage and granulocytic MDSC identities in the TME, MEKi+STAT3i treatment resulted in transcriptional *reprogramming* of the tumor-infiltrating monocytic/macrophage sub-cluster from an alternatively activated M2-like inflammatory phenotype—characterized by enrichment of *Arg1*, *Thbs1*, and *Chil3*—to a classically activated M1-like phenotype—characterized by enrichment of MHC-II genes H2-Eb1, H2-Ab1, and M1-master regulator *Ciita* (*31*) (**Fig. 3D**).

**Figure 3.**
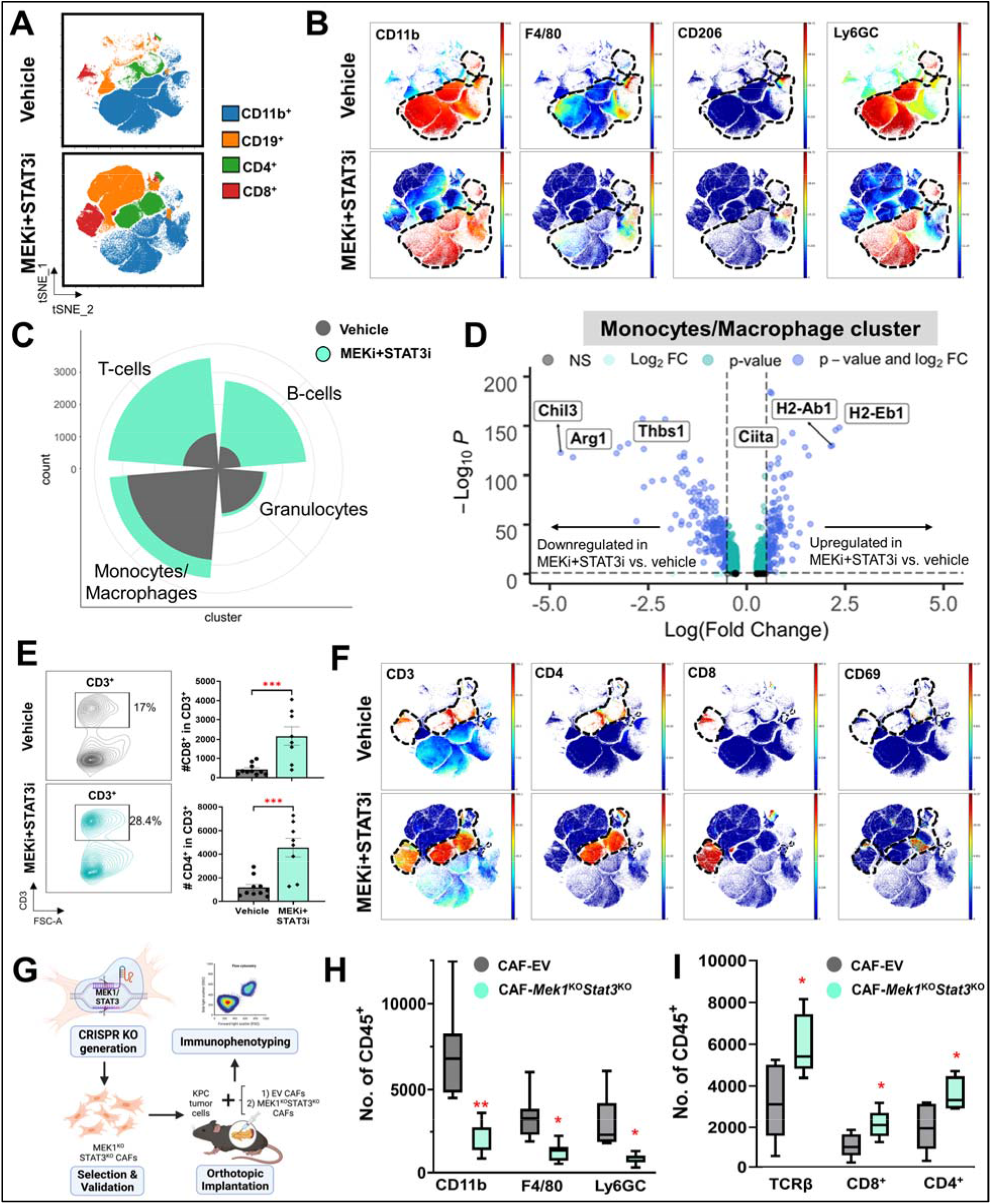
Combined MEK and STAT3 inhibition reprograms the immunosuppressive myeloid microenvironment and facilitates intratumoral T-cell trafficking in part via a CAF-dependent manner. **(A)** Mass cytometry time-of-flight (CyTOF) FlowSOM plots depicting changes in total myeloid (CD11b^+^), T-cell (CD4^+^ and CD8^+^) and B-cell (CD19^+^) populations in PKT mice treated with vehicle or MEKi+STAT3i (n=7-8 mice/arm); **(B)** Representative viSNE plots demonstrating changes in CD11b^+^ myeloid sub-populations F4/80^+^ macrophages, CD206^+^ M2-like macrophages, and Ly6GC myeloid-derived suppressor cells between vehicle- and MEKi+STAT3i-treated PKT mice as analyzed by CyTOF. Parent CD11b^+^ cell populations are denoted by dashed line; **(C)** Circumplex plot from single-cell RNA sequencing (scRNAseq) analysis in PKT mice treated with either vehicle (grey) or MEKi+STAT3i (green) showing relative abundance of T-cell, B-cell, granulocytes, and monocyte/macrophage immune cell clusters between treatment groups; **(D)** Volcano plot of transcripts associated with monocyte/macrophage polarization toward M1 (*Ciita*, *H2-Ab1*, *H2-Eb1*) or M2 (*Chil3*, *Arg1*, *Thbs1*) skewness that are significantly overexpressed (right) or underexpressed (left) in MEKi+STAT3i-treated vs. vehicle-treated scRNAseq monocyte/macrophage transcriptomes. FDR-corrected P-value and log_2_(fold change) thresholds were established at ≤0.05 and ≥0.5, respectively; **(E)** Flow cytometric analysis comparing global CD3^+^ T-cell populations, shown in representative contour plots (*left*), as well as CD4^+^ and CD8^+^ T-cell subsets, shown in histograms (*right*), between vehicle and MEKi+STAT3i-treated PKT mice (n=8-10 mice/arm); **(F)** Representative viSNE plots demonstrating changes in CD3^+^, CD4^+^, CD8^+^, and CD69^+^ T-cell populations between vehicle- and MEKi+STAT3i-treated PKT mice as analyzed by CyTOF. Parent CD3^+^, CD4^+^, or CD8^+^ T-cell populations are denoted by dashed line, where applicable; **(G)** Schematic showing generation of CRISPR/Cas9 genetic silencing of *Mek1* and *Stat3* in KPC cancer-associated fibroblasts, and orthotopic injection of KPC tumor cells with either empty vector (EV) CAFs or *Mek1*^KO^*Stat3*^KO^ CAFs in syngeneic C57B/l6 mice, followed by immunophenotyping of established tumors by flow cytometry. Histograms showing total numbers of **(H)** CD11b^+^, F4/80^+^, and Ly6GC^+^ myeloid cells, and **(I)** TCR-β, CD8^+^, and CD4^+^ T-cells in CAF-EV vs. CAF-*Mek1*^KO^*Stat3*^KO^ tumors (n=7-8 mice/group). Data are shown as mean ± SEM. ns, not significant; *, p<0.05; **, p<0.01; ***, p<0.001.

This remodeling of the innate immune microenvironment following MEKi+STAT3i treatment was associated with significant increases in the proportion of total (CD3^+^), T-helper (CD4^+^), cytotoxic (CD8^+^), and activated (CD69^+^) T-cell populations by flow cytometry (**Fig. 3E**) and CyTOF immunoprofiling (**Fig. 3F**) in PKT mice compared with vehicle treatment. MEKi+STAT3i treatment also improved effector (CD44^+^) and central (CD62L^+^) memory T-cell markers, as well as expanded TCRβ expression, in the PDAC TME compared with vehicle treatment via CyTOF profiling (**Fig. S7B**).

Next, to determine if CAF-restricted silencing of *Mek1/Stat3* recapitulates the immunologic remodeling following systemic MEKi+STAT3i treatment, we performed flow cytometric immunophenotyping in orthotopic models where KPC6694c2 tumor cells were co-injected with either CAF-EV or CAF-*Mek1*^KO^*Stat3*^KO^ (1:9 ratio) in syngeneic C57Bl/6 mice (**Fig. 3G**). Significant reduction in CD11b^+^, F4/80^+^, and Ly6GC^+^ myeloid populations were observed in CAF-*Mek1*^KO^*Stat3*^KO^ compared with CAF-EV tumors (**Fig. 3H**). Conversely, CAF-*Mek1*^KO^*Stat3*^KO^ tumors demonstrated significantly increased TCR-β^+^, CD8^+^, and CD4^+^ T-cell populations compared with CAF-EV tumors (**Fig. 3I**). Taken together, these data suggest that combined MEKi+STAT3i reprograms stromal inflammation to promote an anti-tumor immune microenvironment by decreasing immunosuppressive myeloid cell populations and augmenting T-cell trafficking, in part, via a CAF-dependent mechanism.

### Combined MEKi+STAT3i controls PDAC growth in a T-cell dependent manner *in vivo*

We have previously shown that MEKi+STAT3i results in improved survival compared with vehicle treatment or MEKi or STAT3i monotherapy in PKT mice (*22*). Given the aforementioned immunologic changes with MEKi+STAT3i, we sought to determine if the anti-tumor effects of this combination therapy are T-cell dependent by performing CD4^+^ and CD8^+^ T cell depletion in this model concurrently with MEKi+STAT3i (**Fig. S8A-C**). Non-T-cell depleted PKT mice treated with MEKi+STAT3i showed a significant decrease in tumor weight compared with vehicle treatment and demonstrated 100% survival up to 80 days of treatment (**Fig. 4A&B**). In contrast, T-cell depleted mice treated with MEKi+STAT3i showed no significant decrease in tumor weight compared with vehicle-treated mice (**Fig. 4A**), despite confirmation of pERK and pSTAT3 inhibition in tumor lysates (**Fig. 4C**). Furthermore, the median survival of the T-cell depleted MEKi+STAT3i mice was significantly diminished compared with MEKi+STAT3i treatment without T-cell depletion (median 60.5 vs. not reached [NR] days, respectively; log-rank p=0.0001; **Fig. 4B**). These results show that the decreased tumor burden and improved survival with MEKi+STAT3i in PDAC are T-cell dependent.

**Figure 4.**
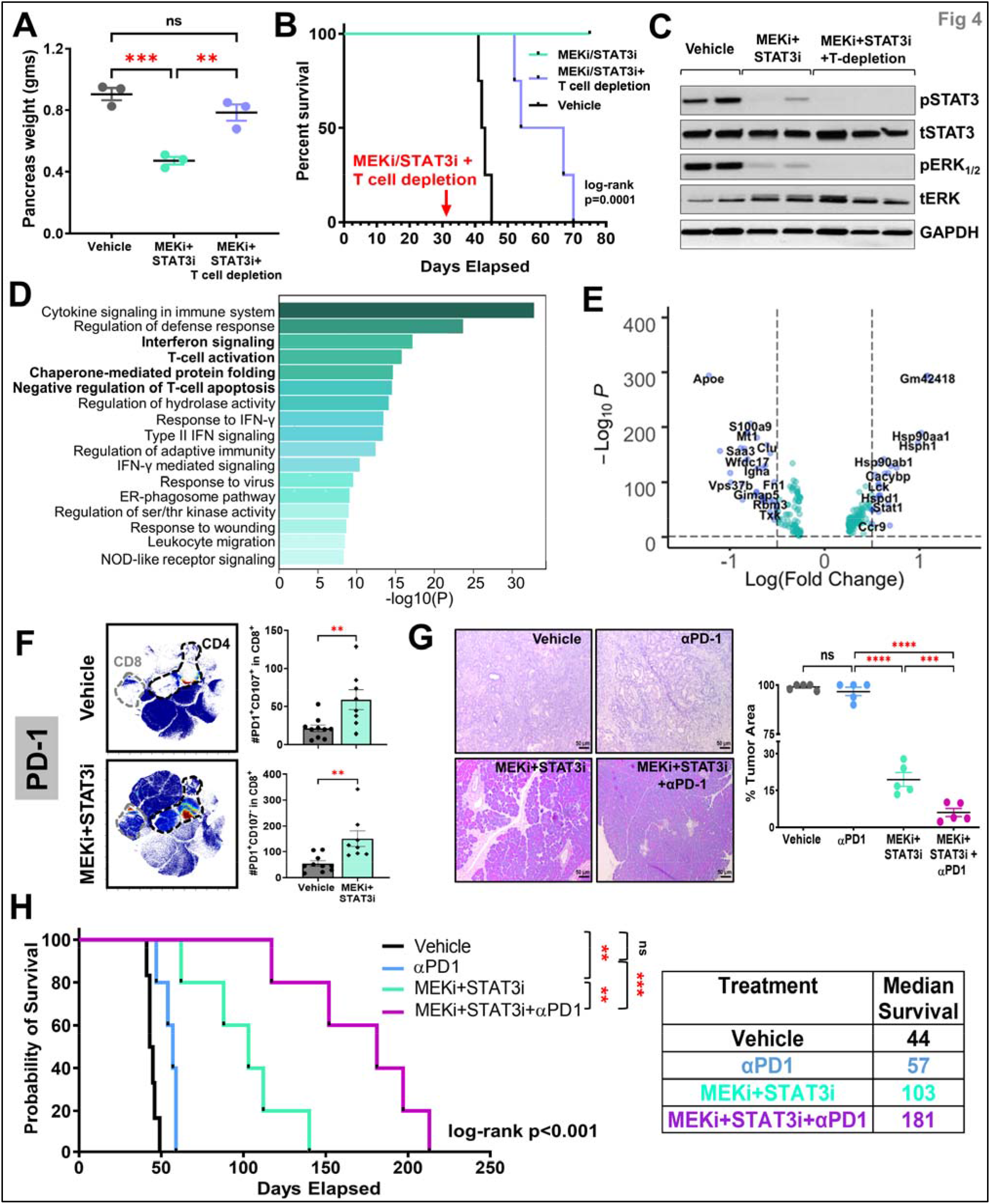
Combined MEK and STAT3 controls PDAC growth in a T-cell dependent manner and overcomes immunotherapy resistance *in vivo*. PKT mice were treated with vehicle, MEKi+STAT3i, or MEKi+STAT3i following T-cell depletion with anti-CD4 and anti-CD8 antibodies (scheme in **Fig. S8**). Differences in **(A)** pancreas weight at sacrifice and **(B)** overall survival were compared among treatment arms. **(C)** Representative western blot demonstrating target inhibition of pERK1/2 and pSTAT3 in mice treated with MEKi+STAT3i ± T-cell depletion; **(D)** Metascape pathway enrichment analysis depicting top 17 signaling pathways from MSigDB compendium differentially upregulated in MEKi+STAT3i-treated compared with vehicle-treated T-cell single-cell transcriptomes from scRNAseq analysis in PKT tumors. Bolded pathways highlight those related to interferon signaling, T-cell activation, chaperone-mediated protein folding, and negative regulation of T-cell apoptosis; **(E)** Volcano plot of transcripts associated with aforementioned pathways that are significantly overexpressed (right) or underexpressed (left) in MEKi+STAT3i-treated vs. vehicle-treated single cell T-cell transcriptomes. FDR-corrected P-value and log2(fold change) thresholds were established at ≤0.05 and ≥0.5, respectively; **(F)** Mass cytometry time-of-flight (CyTOF) analysis of PKT mice treated with vehicle or MEKi+STAT3i showing an increase in PD-1^+^ tumor-infiltrating CD8^+^ T-cells, as visualized by representative viSNE plots (**F**, *left*). Flow cytometric analysis showing increase in degranulating CD8^+^PD-1^+^CD107a^+^ T-cells and non-degranulating CD8^+^PD-1^+^CD107a^-^ T-cells in MEKi+STAT3i treated vs. vehicle-treated mice (n=8-10 mice/arm) (**F**, *right*); **(G)** Representative images from H&E sections from PKT mice treated with vehicle, αPD-1, MEKi+STAT3i, or MEKi+STAT3i plus αPD-1 for 4 weeks (**G**, *left*). Percent tumor area (**G**, *right*) at endpoint sacrifice were compared between treatment arms (n=5 mice/arm); **(H)** Kaplan-Meier survival plot depicting overall survival in PKT mice treated with vehicle, αPD-1, MEKi+STAT3i, or MEKi+STAT3i plus αPD-1 beginning at 4-4.5 weeks of age. Log-rank survival comparisons between individual treatment groups and overall cohort, as well as adjoining table with median survival times, are provided. Where applicable, data are shown as mean ± SEM. ns, not significant; *, p<0.05; **, p<0.01; ***, p<0.001.

### MEKi+STAT3i treatment promotes a distinct transcriptional and functional reprogramming in tumor-infiltrating T-cells

We next sought to investigate if transcriptional and functional reprogramming of the T-cell compartment underlies the T-cell dependent effects of MEKi+STAT3i-induced tumor control. Interrogation of single-cell T-cell transcriptomes revealed global skewness towards effector versus exhausted programming in the T-cell compartment of MEKi/STAT3i- vs. vehicle-treated tumors (**Fig. S9A**). In particular, gene ontology analysis of T-cell transcriptomes in MEKi+STAT3i-treated tumors revealed enrichment of pathways related to interferon signaling, T-cell activation, chaperone-mediated protein folding, and negative regulation of T-cell apoptosis (**Fig. 4D**). In particular, we observed differential regulation of genes related to TCR activation (e.g., lymphocyte-specific protein tyrosine kinase *Lck* upregulation (*32*)), anti-exhaustion programming (e.g., *Stat1* upregulation (*33*)*, Apoe* downregulation), effector differentiation and function (e.g., heat-shock chaperone protein *Hsp90aa1, Hsp90ab1, Hspd1, Hsph1* upregulation (*34*)) in T-cell transcriptomes in MEKi+STAT3i- vs. vehicle-treated PKT tumors (**Fig. 4E**).

To further explore the functional status of tumor-infiltrating CD8^+^ T-cells in MEKi+STAT3i- vs. vehicle-treated PKT mice, we examined CD8^+^ T-cell antigen experience (PD1^+^) and degranulating activity (CD107a^+^) in these cells. CyTOF profiling revealed a significant upregulation of PD1^+^ tumor-infiltrating CD8^+^ T-cells in PKT tumors treated with MEKi+STAT3i compared with vehicle treatment (**Fig. 4F** *left*). Despite a significant increase in antigen-experienced degranulating CD8^+^PD1^+^CD107a^+^ T-cells following MEKi+STAT3i, we also observed an increased proportion of PD1^+^CD107a^-^ non-degranulating CD8^+^ T-cells (**Fig. 4F** *right*). Of note, we found no significant changes in PD-L1 expression in PKT tumors between vehicle treatment, MEKi or STAT3i monotherapy, and combined MEKi+STAT3i treatment (**Fig. S9B&C**). Taken together, these data suggest that MEKi+STAT3i treatment—directly or indirectly—may uncouple effector/anti-apoptotic transcriptional programs from dysfunctional programs in tumor-infiltrating CD8^+^ T-cells to augment their degranulating capacity *in vivo*.

### Treatment with MEKi+STAT3i overcomes resistance to PD-1 blockade in PKT mice

Viewed broadly, treatment with MEKi+STAT3i addresses the key barriers to ICI sensitivity in PDAC: stromal inflammation and IL-6/CXCL1-generating iCAFs, immunotherapy-resistant *Lrrc15*^+^ myCAF populations, an immunosuppressive myeloid cell-dominant TME, and T-cell exclusion and dysfunctional programming. These data suggest that the addition of MEKi+STAT3i to ICI may overcome resistance to immunotherapy in immunologically inert PDAC. To test this hypothesis, PKT mice were treated with vehicle, αPD-1 monotherapy, MEKi+STAT3i, and MEKi+STAT3i combined with αPD-1 antibody (**Fig. S10A**). Consistent with our previous data (*22*), MEKi+STAT3i significantly reduced tumor burden compared with vehicle treatment in PKT mice. Moreover, in keeping with prior evidence in KPC mice (*35*) as well as clinical trials of ICI in PDAC patients (*5, 6*), αPD-1 monotherapy had no effect on tumor burden compared with vehicle treatment in PKT mice, further validating the relevance of our model as ICI-resistant. Importantly, the addition of MEKi+STAT3i to αPD-1 blockade significantly reduced tumor burden (**Fig. 4G**) and pancreas weight (**Fig. S10B**) compared to αPD-1-treated as well as MEKi+STAT3i-treated mice. In fact, in this aggressive PKT PDAC GEMM, where the pancreata of vehicle- and αPD-1-treated mice are completely replaced by tumor by 6 weeks of age, MEKi+STAT3i treated mice show only 19% tumor area, on average, after 4 weeks of treatment (∼8 weeks of age), while the addition of MEKi+STAT3i to αPD-1 treatment maintained normal pancreas architecture with <6% tumor area (**Fig. 4G** *left*).

### Addition of MEKi+STAT3i to PD-1 blockade significantly improves survival in PKT mice

We next sought to determine if this combination therapy overcomes resistance to ICI and improves survival in PKT mice (treatment schema in **Fig. S10C**). Confirming our previous data, MEKi+STAT3i significantly increased survival compared with vehicle treatment (median 103 vs. 44 days; p=0.0013). Consistent with the lack of response to αPD-1 monotherapy in PDAC GEMMs (*36–39*), we observed no difference in survival between αPD-1 monotherapy and vehicle treatment (median 57 vs. 44 days, p=0.34). Importantly, the addition of MEKi+STAT3i to PD-1 blockade dramatically improved PKT survival compared with αPD-1 monotherapy (median 181 vs. 57 days, p=0.0005), as well as combined MEKi+STAT3i (median 181 vs. 103 days, p=0.0064; **Fig. 4H**). Furthermore, there was no difference in body weight between vehicle and treatment groups, indicating that the combination of MEKi+STAT3i with PD-1 blockade is well tolerated with no additional toxicity compared with either αPD-1 monotherapy or MEKi+STAT3i *in vivo* (**Fig. S10D**). These results highlight the significant role of MEKi+STAT3i in overcoming resistance to ICI in an immunologically inert autochthonous preclinical model of PDAC.

### The addition of MEKi+STAT3i to PD-1 blockade improves recruitment, activation, and functional cytotoxicity of tumor-infiltrating T-cells

Next, we examined the immunologic underpinnings of the dramatic improvement in survival of PKT mice treated with combined MEKi+STAT3i and αPD-1 compared with αPD-1 monotherapy. CyTOF-based immune profiling demonstrated that, compared with both vehicle treatment and αPD-1 monotherapy, the addition of MEKi+STAT3i to PD-1 blockade resulted in a significant increase in tumor-infiltrating CD4^+^ and CD8^+^ T-cells, as well as CD3^+^ T-cells demonstrating T-cell activation markers CD44 and CD69 and central memory marker CD62L (**Fig. 5A**). These broad-based changes in adaptive immune populations were then corroborated by flow cytometry in the CD4^+^ and CD8^+^ T-cell compartments separately. In both CD4^+^ and CD8^+^ T-cell compartments alike, compared to αPD-1 monotherapy or vehicle treatment, the addition of MEKi+STAT3i to PD-1 blockade significantly augmented proportions of activated (CD69^+^), effector memory (CD44^+^CD62L^-^), central memory (CD44^+^CD62L^+^), tissue resident memory (CD44^+^CD62L^-^CD103^+^), and degranulating effector (CD44^+^CD62L^-^CD107a^+^) tumor-infiltrating T-cells (**Fig. 5B, Fig. S11**).

**Figure 5.**
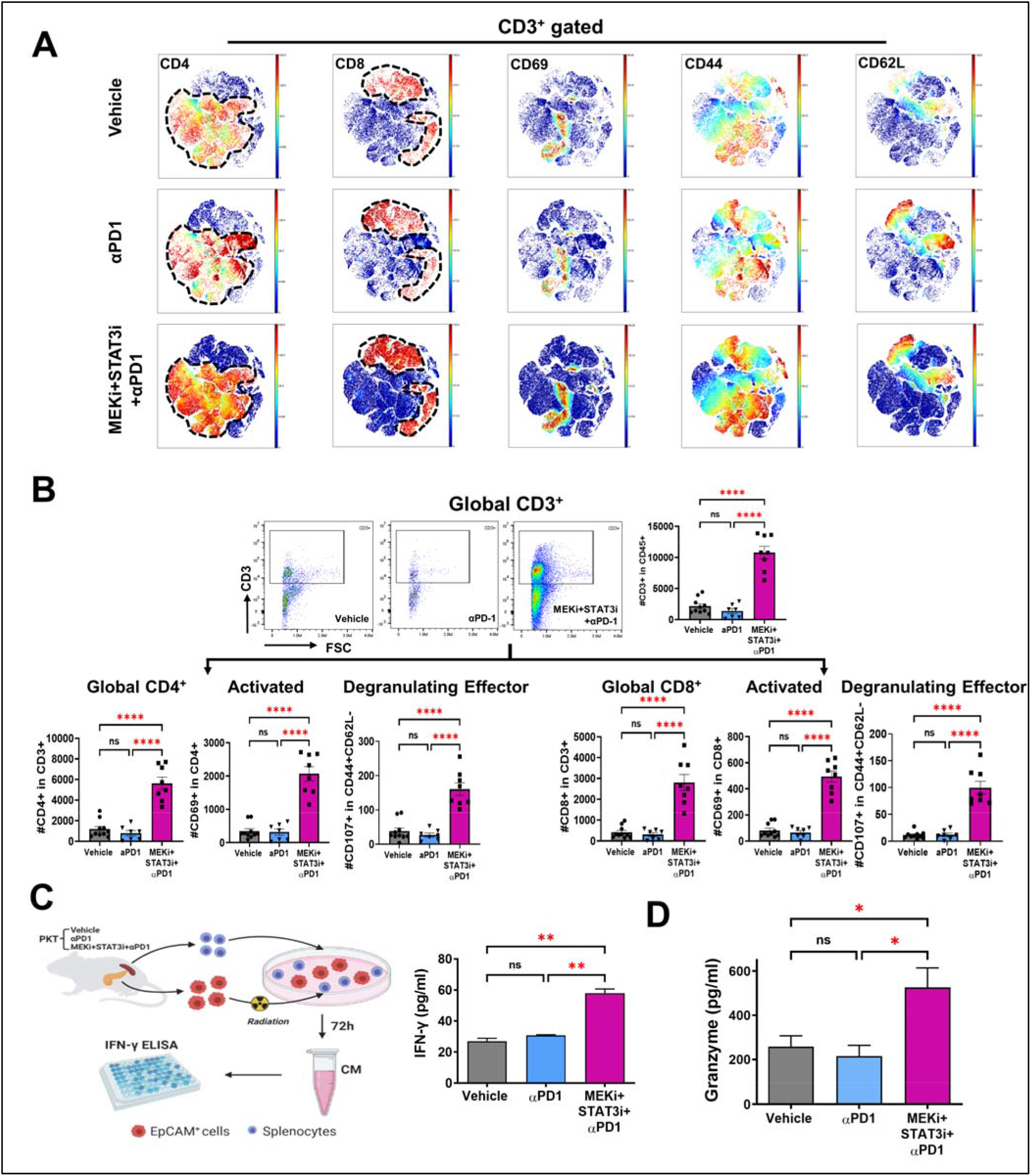
Addition of MEKi+STAT3i to PD-1 blockade augments recruitment, activation, and functional cytotoxicity of tumor-infiltrating T-cells. **(A)** viSNE plots demonstrating levels of tumor-infiltrating CD3^+^ cell subsets (CD4^+^, CD8^+^, CD69^+^, CD44^+^, and CD62L^+^) in vehicle, αPD-1, and combined MEKi+STAT3i/αPD-1 treated PKT mice as analyzed by CyTOF (n=7-8 mice/arm); **(B)** Analysis by flow cytometry showing changes in total CD3^+^, CD4^+^, and CD8^+^ tumor-infiltrating T-cells as well as levels of activated CD69^+^ and degranulating effector (CD4^+^CD62^-^CD107^+^) CD4^+^/CD8^+^ T-cells in PKT mice treated with vehicle, αPD-1, or MEKi+STAT3i/αPD-1 (n=8-10 mice/arm); **(C)** Schematic depicting *ex vivo* tumor cell:splenocyte co-culture experiment. Splenocytes were isolated from PKT mice following treatment with vehicle, αPD-1, and MEKi+STAT3i/αPD-1 and co-cultured with irradiated PKT tumor cells for 72 hours. IFN-γ release was determined by ELISA (right). **(D)** ELISA demonstrating granzyme B levels in total tumor lysate in vehicle, αPD1, and MEKi+STAT3i/αPD-1 treatment arms (n=3 mice/arm). Data are shown as mean±SEM. Scale bar = 50 µm. ns, not significant; *, p<0.05; **, p<0.01; ***, p<0.001; ****, p<0.0001.

Given significant improvements in the recruitment and activation profile of tumor-infiltrating T-cells, we next investigated if the addition of MEKi+STAT3i to PD-1 blockade enhances the functional cytotoxicity of CD8^+^ T-cells compared with αPD-1 monotherapy and vehicle treatment. Using *ex vivo* CD8^+^ T-cell:tumor cell IFN-γ release assays, PKT-derived EpCAM^+^ tumor cells were co-cultured with splenocyte-derived CD8^+^ T-cells from PKT mice treated with vehicle, αPD-1 alone and MEKi+STAT3i/αPD-1, and the co-culture conditioned media subjected to IFN-γ ELISA. The addition of MEKi+STAT3i to PD-1 blockade significantly increased the cytolytic activity of CD8^+^ T-cells compared with αPD-1 monotherapy or vehicle treatment (**Fig. 5C**). Further validating these functional T-cell data, the addition of MEKi+STAT3i to PD-1 blockade significantly increased tumor-specific granzyme B levels in PKT tumor lysates, compared with αPD-1 monotherapy or vehicle treatment (**Fig. 5D**). These data collectively show that the addition of MEKi+STAT3i to PD-1 blockade invigorates the adaptive immune compartment to overcome T-cell dysfunction and ICI resistance in the PDAC TME. Moreover, it is plausible that MEKi+STAT3i, which improves tumor-infiltrating CD8^+^ T-cell antigen experience but does not completely rescue their degranulating capacity (**Fig. 4F**), primes an activated CD8^+^ T-cell compartment for ICI to further augment the cytolytic function of PDAC-infiltrating CD8^+^ T-cells.

### Dampening of suppressive innate immune microenvironment in PKT mice is driven by MEKi+STAT3i but not anti-PD-1 monotherapy

Given recent evidence that myeloid cell-specific PD-1 ablation—independent of T-cell-directed anti-PD-1 effects—may contribute to effector T-cell infiltration and antitumor immunity (*40*), we sought to investigate the differential effects of PD-1 blockade alone and combined MEKi+STAT3i/αPD-1 on the innate immune microenvironment in PKT tumors. CyTOF-based immune profiling revealed that αPD-1 monotherapy had no impact on innate immune infiltration in PKT tumors compared with vehicle treatment. The addition of MEKi+STAT3i to PD-1 blockade, however, significantly reduced the recruitment of tumor-infiltrating F4/80^+^ macrophages, CD206^+^ M2-like macrophages, and Ly6GC^+^ MDSCs compared with αPD-1 monotherapy or vehicle treatment (**Fig. 6A**). Flow cytometric analysis (**Fig. 6B**) and tissue-based immunofluorescence (**Fig. 6C**) confirmed a significant decrease in CD11b^+^ myeloid, F4/80^+^ macrophages, and Ly6G^+^ MDSCs following MEKi+STAT3i/αPD-1 compared with αPD-1 monotherapy or vehicle treatment. These data reinforce that the dampening of immunosuppressive innate populations in the PDAC TME is driven predominantly by MEKi+STAT3i without significant contributions from αPD-1 monotherapy, and may underlie the adaptive immune invigoration observed with combination MEKi+STAT3i and PD-1 blockade.

**Figure 6:**
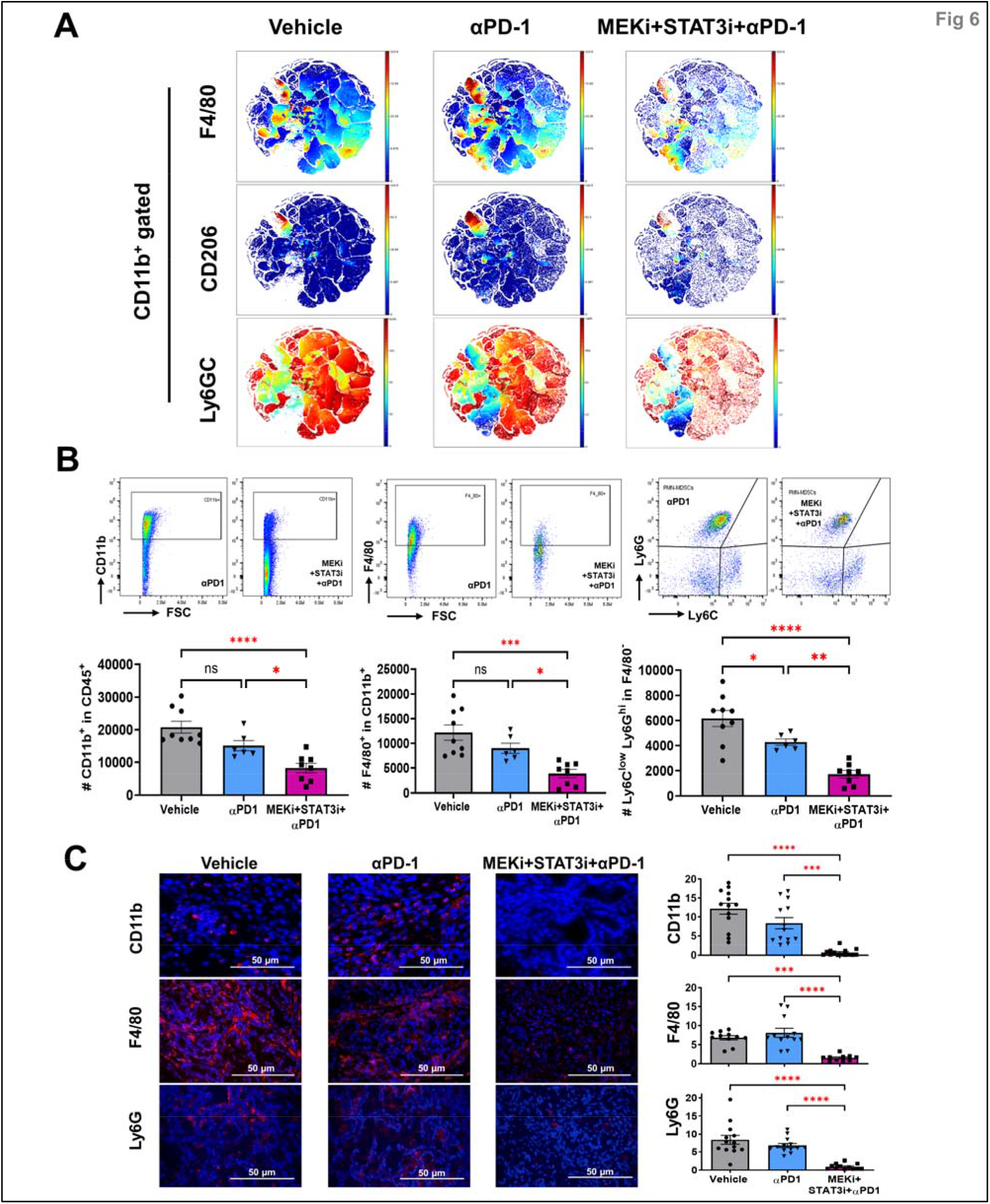
Differential effects on innate immune remodeling are driven by MEKi/STAT3i with minimal contribution from anti-PD-1 monotherapy in PKT mice. (**A**) viSNE plots depicting mass cytometry time-of-flight (CyTOF) analysis of CD11b^+^ myeloid cell subsets (F4/80^+^, CD206^+^, and Ly6G^+^C^+^) in PKT mice treated with vehicle, αPD-1, and MEKi+STAT3i plus αPD-1 for 4 weeks (n=7-8 mice/arm); **(B)** Levels of total myeloid (CD11b^+^), macrophage (CD11b^+^F4/80^+^), and PMN-MDSCs (Ly6G^+^Ly6C^lo^F4/80^-^) cells were determined by flow cytometry in PKT tumors treated with vehicle, αPD1, and MEKi+STAT3i+αPD1 (n=7-8 mice/arm); **(C)** Immunofluorescent staining and quantification of CD11b^+^, F4/80^+^, and Ly6G^+^ levels in PKT mice among indicated treatment arms vehicle, αPD-1, and MEKi+STAT3i plus αPD-1 (n=3-4 mice/arm). Data are shown as mean ± SEM. Scale bar = 50 µm. ns, not significant; *, p<0.05; **, p<0.01; ***, p<0.001; ****, p<0.0001.

### Combination MEKi+STAT3i and PD-1 blockade demonstrates efficacy in a metastatic PDAC patient with chemotherapy-refractory disease

A 69-year-old male with a remote history of Hodgkin’s lymphoma presented with a resectable microsatellite stable, low-tumor mutation burden (TMB) PDAC in the pancreatic body. He received neoadjuvant fluorouracil (5-FU), leucovorin, irinotecan, and oxaliplatin (FOLFIRINOX) for three months before undergoing curative-intent distal pancreatectomy and splenectomy. Pathology revealed a margin-negative resection, and a 2-cm focus of residual adenocarcinoma with 0/33 involved lymph nodes. He then completed three months of adjuvant FOLFIRINOX (**Fig. 7A**). Two months following completion of chemotherapy, he was diagnosed with a local recurrence in the pancreatic bed as well as a metastatic lesion in segment VII of the liver (**Fig. 7B**). He initiated second-line chemotherapy with gemcitabine/*nab*-paclitaxel but had progression of disease with a significant reduction in performance status.

**Figure 7:**
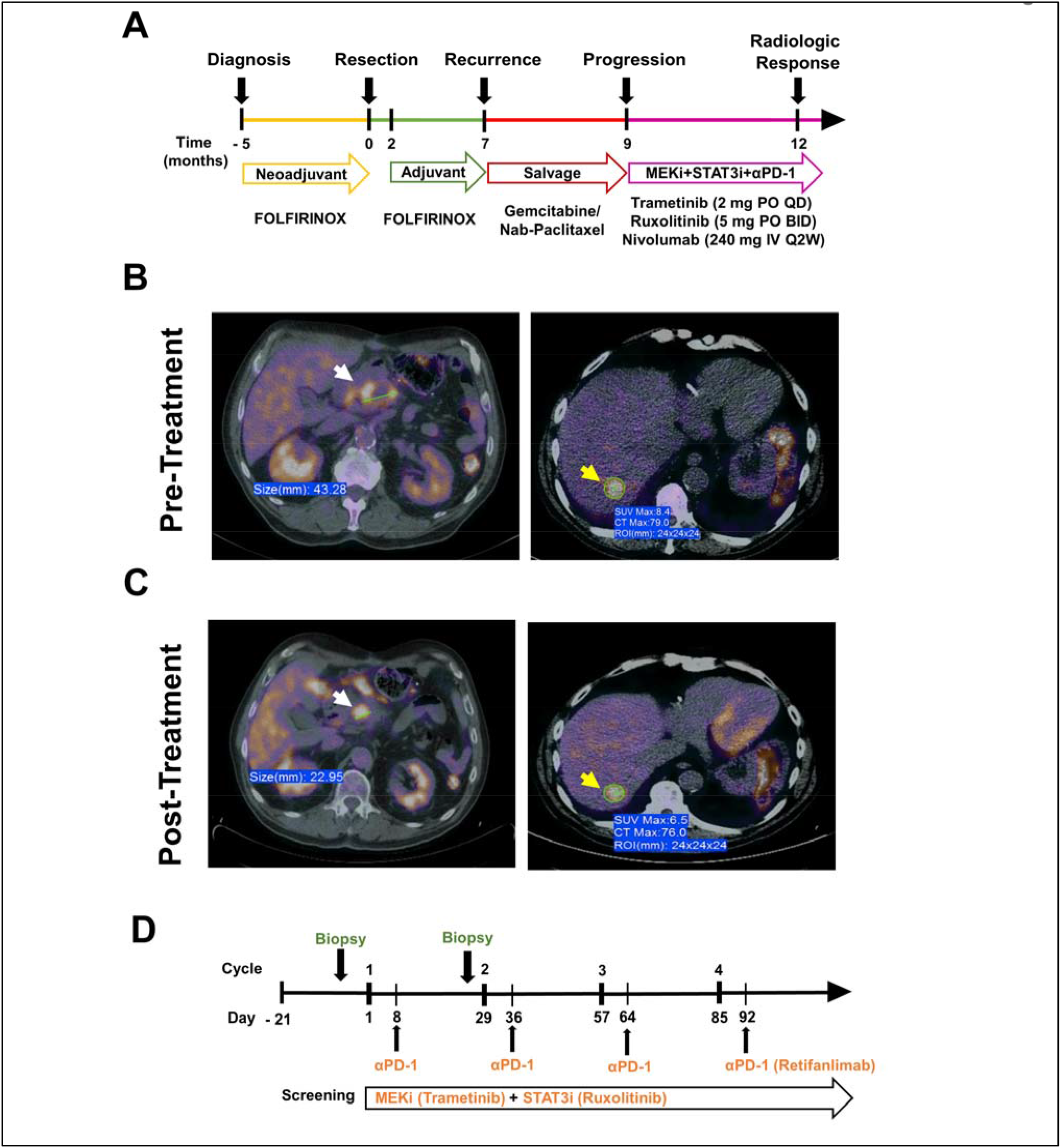
Combination MEK, STAT3, and PD-1 inhibition demonstrates efficacy in a metastatic PDAC patient with chemotherapy-refractory disease, laying groundwork for clinical trial. **(A)** Treatment timeline for patient with chemotherapy refractory PDAC prior to initiation of Trametinib, Ruxolitinib, and Nivolumab treatment. Pre-treatment **(B)** and post-treatment **(C)** PET/CT scan showing a significant reduction in size and FDG avidity of both locally recurrent tumor in pancreatic bed (white arrows) and in a segment VII liver metastasis (yellow arrows) on PET/CT imaging following 3 months of treatment with Trametinib, Ruxolitinib, and Nivolumab; **(D)** Design of Phase 1 clinical trial investigating Trametinib (MEKi), Ruxolitinib (STAT3i), and Retifanlimab (αPD-1) in advanced/metastatic pancreatic cancer patients. Proposed treatment schedules, as well as scheme for biopsies/blood collection for correlative endpoints, are shown.

After multidisciplinary discussion, he received off-label combination treatment with MEKi Trametinib (Mekinist/Novartis, 2mg PO QD), STAT3i Ruxolitinib (Jakafi/Incyte, 5mg PO BID), and αPD-1 Nivolumab (Opdivo/Bristol-Myers Squibb, 240mg IV Q2W) (**Fig. 7A**). Within one month of commencing treatment, the patient experienced a dramatic improvement in clinical and performance status. Follow-up PET/CT obtained after 3 months of treatment demonstrated significant response in the pancreatic bed mass, with nearly 50% reduction in size from 43 mm to 23 mm and a metabolic response with reduction in SUV_max_ from 8.4 to 6.5 post-treatment in the liver metastasis on PET/CT (**Fig. 7C**). The regimen was well-tolerated with no reported major adverse events while on therapy, and resulted in significant improvement in the patient’s performance status and self-reported quality-of-life.

Based on extensive preclinical rationale detailed here, as well as encouraging signals of its clinical efficacy and tolerability, we are pursuing an investigator-initiated clinical trial investigating this treatment combination in advanced PDAC patients. Briefly, patients with metastatic PDAC who are refractory to first-line standard therapy will be treated in a phase 1 trial with trametinib (MEKi), ruxolitinib (STAT3i) and retifanlimab (anti-PD-1 antibody) to determine the safety of the combination and the recommended dose/schedule for phase 2 development. Pre- and post-treatment blood and tumor specimens will be collected; RNA-sequencing and multiparametric CyTOF profiling in peripheral blood samples will be used to identify novel transcriptomic and immunologic signatures and circulating immune cellular phenotypes of response following immunomodulatory MEKi+STAT3i/α-PD1 treatment (**Fig. 7D**).

## DISCUSSION

The present manuscript uncovers a novel treatment strategy that remodels stromal inflammation and myeloid-enriched immune microenvironments to activate functional T-cell anti-tumor immunity and overcome ICI resistance in PDAC. Resistance to ICI is a hallmark of PDAC (*5, 6*), defined by T-cell exclusion and tissue-level immune tolerance driven by *KRAS*-mediated tumor-intrinsic programs (*1, 20*) as well as interwoven suppressive cellular networks comprising innate immune populations and tumor-permissive CAF signaling (*7*). Our therapeutic approach using MEKi+STAT3i highlighted here not only disrupts oncogenic signaling via reciprocally activated RAS/MEK/ERK and JAK/STAT3 pathways in PDAC tumor cells (*22*) but also addresses the critical barriers to ICI resistance inherent in the PDAC TME: stromal inflammation via IL-6/CXCL1-secreting iCAFs (*12, 14, 15*), *Lrrc15*^+^ myCAF-driven immunotherapy resistance (*9*), immunosuppressive myeloid cell-enriched tumor ecosystems, and T-cell exclusion and dysfunctional programming (*3, 7*). We leverage these mechanistic underpinnings in a highly aggressive and ICI-resistant preclinical model to demonstrate an impressive improvement in tumor burden and survival when combining MEKi+STAT3i with PD-1 inhibition, compared with either MEKi+STAT3i alone or anti-PD1 monotherapy. Importantly, the efficacy and tolerability of Trametinib (MEKi), Ruxolitinib (STAT3i), and Nivolumab (anti-PD1) in a patient with metastatic chemo-refractory PDAC offers encouraging signals of its clinical translatability, and is currently being pursued in an investigator-initiated phase 1 trial.

Emerging evidence has elucidated the signaling crosstalk between tumor cells and CAFs in promoting hallmarks of its aggressive biology, such as epithelial-to-mesenchymal transition and proliferative capacity (*41*). In addition to our group’s previous discovery of reciprocally activated MEK/ERK (downstream of RAS) and STAT3 signaling in tumor cells promoting therapeutic resistance (*22, 23*), other groups have reported CAF-mediated factors such as TGF-β in further instigating tumor cell-specific MEK/ERK and STAT3 co-activation to perpetuate such therapy-resistant signaling (*41*). It is plausible, therefore, that combined MEKi+STAT3i treatment restrains PDAC outgrowth by not only ameliorating reciprocal MEK/ERK and STAT3 signaling in tumor cells, but also dampening tumor-permissive tumor-stromal crosstalk via attenuation of TGF-β^+^ myofibroblastic CAF populations in the PDAC TME (see **Fig. 2**).

In parallel with these observations, our scRNAseq analyses demonstrate emergence of a mesenchymal stem/stromal cell (MSC) progenitor-like CAF subset in MEKi+STAT3i-treated PKT tumors, defined by expression of MSC markers *Cd34*, *Ly6a*, *Ly6c1*, and Meflin (*Lslr*). A recent study from Mizutani and colleagues revealed that Meflin^+^ MSC-CAFs were precursors to α-SMA^+^ myCAF populations—consistent with our pseudotime lineage reconstruction studies (see **Fig. 2G**)—and that genetic ablation of Meflin^+^ MSC-CAFs resulted in significant tumor progression with poorly differentiated histology in a PDAC model (*28*), suggesting that Meflin^+^ MSC-CAF subtypes may be tumor-restraining. As such, a more provocative mechanism by which MEKi+STAT3i exerts tumor control may be by uncovering stromal plasticity and reprogramming the CAF compartment towards a putative *tumor-restraining* MSC-like fibroblast phenotype. Ongoing lineage tracing studies in our laboratory are deciphering whether reciprocal co-activation of MEK and STAT3 signaling—operative in the tumor-cell compartment—also drives mesenchymal progenitor-like CAF progression to myofibroblastic CAF phenotypes in PDAC. It is also possible that these MSC-like CAFs emerge from an alternative fate trajectory of mesenchymal progenitors in the absence of TGFβ^+^ myCAF signaling following MEKi+STAT3i treatment.

This reprogramming of the myofibroblastic CAF compartment may also underlie MEKi+STAT3i-induced sensitization to ICI in our preclinical models. Growing evidence highlights a prominent role for CAFs in restraining efficacy of ICI in multiple solid tumors (*42, 43*), and implicates subtype-specific heterogeneity in myCAF populations as a primary driver of immunotherapy resistance (*9, 44*). For instance, myCAF subpopulations in breast cancer patients nominated as ecm-myCAF (overexpressing *LRRC15* and *GBJ2*) and TGFβ-myCAF (overexpressing *CST1* and *TGF*β*1*) govern ICI sensitivity via a regulatory T-cell-dependent mechanism (*44*). Consistent with these findings, Dominguez and colleagues demonstrated that an *Lrrc15*^+^ CAF signature is a marker of TGF-β-driven myCAFs in the PDAC TME, is abundantly distributed in human PDAC specimens, and underlies poor response to immunotherapy in multiple solid cancers (*9*). As such, one of the major mechanisms by which MEKi+STAT3i overcomes immunotherapy resistance in the present study may be by attenuating immunosuppressive *Lrrc15*^+^ myCAF sub-phenotypes in the PDAC TME (see **Fig. S5E**). In our upcoming phase 1 clinical trial, we will interrogate the dynamics of stromally-expressed LRRC15 during treatment and its potential as a biomarker of response to MEKi+STAT3i/anti-PD1 therapy.

The present study adds to the growing taxonomy and ontogeny of heterogenous CAF subtypes in the PDAC TME by describing the emergence of a mesenchymal progenitor-like CAF following MEKi+STAT3i, which transcriptionally resembles the FB2-CAF detected in early *Kras*^LSL-G12D^;*Cdkn2a*^flox/flox^;*Ptf1a*^Cre/+^ (KIC) tumors but progressively lost during PDAC progression (*26*). Intriguingly, FB2 CAFs (*Ly6a*^+^*Ly6c1*^+^) appeared to be an incipient progenitor-like state that converged towards *Il6*-expressing FB1 fibroblasts—representing secretory/inflammatory CAFs (*26*). Our pseudotime lineage reconstruction analysis strengthens this observation by suggesting that bone marrow-derived MSC-like fibroblast populations (*Cd34*^+^*Ly6a*^+^*Ly6c1*^+^) may be precursors for mesothelial-derived CAF populations that predominate in the “mature” PDAC TME. As such, one might speculate that a mesenchymal-to-mesothelial transition is an incipient event in CAF lineage commitment—as has been described in developmental fibroblast biology (*45*)—with the first minor trajectory bifurcation yielding *Il6*^+^*Cxcl1*^+^*Il33*^+^ CAF subpopulations with secretory/inflammatory features (see **Fig. 2G**). If this hypothesis is true, treatment with MEKi+STAT3i uncovers stromal plasticity and mitigates stromal inflammation by either: (1) arresting the chronology of this lineage commitment fate from mesenchymal progenitor CAFs to secretory/inflammatory CAFs; and/or (2) promoting dynamic transcriptional reprogramming of secretory/inflammatory CAFs towards its MSC progenitor-like state.

Mechanistically, it appears that MEKi+STAT3i-induced remodeling of stromal inflammation and the innate immune-enriched microenvironment is *in part* CAF-dependent, since genetic silencing of CAF-restricted *Mek1*/*Stat3* not only dampens iCAF polarization but also significantly reduces intratumoral TAM/MDSC infiltration. Beyond these *indirect* CAF-mediated effects on the myeloid landscape, it is also possible that *direct* effects of MEKi+STAT3i treatment on myeloid cellular signaling and/or myelopoiesis contributes to the observed monocyte/macrophage reprogramming from an M2- to M1-like fate in the PDAC TME. While the tolerogenic role of myeloid-specific STAT3 signaling has been well established in pancreatic and other solid tumors (*46*), the role of myeloid-specific MEK/ERK signaling (*47*) and its immunologic repercussions is incompletely understood. The findings herein warrant further investigation into the differential contributions of MEK/ERK and STAT3 signaling in myelopoiesis, myeloid-specific lineage commitment, and multi-directional crosstalk with disparate immune constituents of the PDAC TME.

In summary, our data present novel insight into a combinatorial targeted therapy regimen that overcomes ICI resistance in PDAC by unmasking stromal plasticity and mitigating stromal inflammation, reprogramming the immunosuppressive TME (*3*), and establishing an immunotherapy-permissive T-cell landscape. These data provide novel and timely insight into the concept that context-dependent targeting of inflammatory and/or myofibroblastic CAFs with preservation (or selection) of their tumor-restraining CAF counterparts—and not indiscriminate stromal depletion (*48*)—will be more effective in overcoming immune exclusion and therapeutic resistance in PDAC. Furthermore, the immunologic changes associated with stromal remodeling following MEKi+STAT3i treatment and the T-cell dependence of MEKi+STAT3i-mediated tumor control underscores a complex relationship between diverging signaling cues from heterogenous CAF subtypes, tolerogenic myeloid cell signaling, and effector immune infiltration and programming in the PDAC TME (*49*). Our long-standing efforts in understanding the multi-faceted role of MEK/ERK and JAK/STAT3 signaling in PDAC culminate with clinical evidence of exceptional response to combination MEKi+STAT3i plus PD1 inhibition in a chemorefractory patient with metastatic PDAC. While such dramatic responses are not uniformly expected in all patients enrolled in our upcoming phase 1 trial investigating this novel combination regimen in advanced PDAC patients, the biologic consequences of MEKi+STAT3i treatment *in vivo* described here will be incorporated into a predictive biomarker discovery platform that may inform strategic selection of future patients for these immunotherapeutic combinations.

## MATERIALS AND METHODS

### Study design

The objective of this study was to elucidate the stromal and immunologic mechanisms underlying how MEKi+STAT3i overcomes immunotherapy resistance in PDAC, with the ultimate goal of translating this combinatorial strategy to PDAC patients with treatment-refractory advanced disease. To dissect the impact of MEKi+STAT3i on stromal CAFs *in vivo*, scRNAseq was performed in pooled PKT tumors treated with MEKi+STAT3i or vehicle control (n=3 each) to investigate not only CAF transcriptomic heterogeneity but also CAF evolutionary dynamics uncovered by MEKi+STAT3i treatment at single-cell resolution. Detailed scRNAseq analysis algorithms are described herein. The results of scRNAseq in CAF compartments were validated by whole tumor-derived RNA qPCR in biological triplicates. To investigate the effect of this stromal reprogramming of myeloid and T-cell populations, immune profiling in PKT mice randomized to MEKi+STAT3i or vehicle treatment was performed using TOF mass cytometry and multiparameter flow cytometry. To determine if these MEKi+STAT3i-mediated effects on stromal inflammation and immune remodeling are CAF-dependent, CRISPR/Cas9 genetic editing was performed in PDPN^+^ CAFs isolated from KPC mice to silence *Mek1, Stat3*, or both. Genetic ablation of *Mek1*, *Stat3*, and both *Mek1*/*Stat3* in CAFs were confirmed by western blot, and respective CAF constructs characterized by bulk RNA sequencing compared with empty vector (EV) CAFs in triplicates. CAF-*Mek1*^KO^*Stat3*^KO^ and CAF-EV (90x10^3^ cells/mouse) were co-injected with KPC6694c2 tumor cells (10x10^3^ cells/mouse) orthotopically in syngeneic C57Bl/6 mice pancreata, and tumors at designated endpoint analysis were characterized by multiparameter flow cytometry.

PKT mice enrolled in endpoint and survival studies were randomly assigned (n=8-10 mice/group) to treatment groups vehicle, MEKi+STAT3i, anti-PD1, or MEKi+STAT3i+anti-PD1, and no animals were excluded from the experiment. For *in vivo* studies, the number of animals was selected on the basis of variability observed in pilot experiments and on availability based on genotyping results. Detailed methods are provided in the **Supplementary Materials**, including drug dosages (**Fig. S10**), descriptions of antibodies used for cyTOF, flow cytometry experiments, etc. (**Table S1-S2, S4-S5**) and guide RNAs for CRISPR/cas9 experiments (**Table S3**). All animal experiments were performed in compliance with the ethical guidelines of the Institutional Animal Care and Use Committee at the University of Miami (Protocol #21-057).

A patient with treatment-refractory metastatic PDAC demonstrating progression on two standard lines of systemic chemotherapy was treated off-label with Trametinib (2mg PO QD), Ruxolitinib (5mg PO BID), and Nivolumab (240mg IV Q2W) at the University of Miami Sylvester Comprehensive Cancer Center, and clinical parameters as well as pre- and post-treatment PET/CT images documented.

### In vivo studies

Tumor-bearing PKT mice were generated as previously described (*50*). For endpoint analyses (i.e., flow cytometry, CyTOF, histology), PKT mice were treated with vehicle (0.5% HPMC + 0.1% Tween 80), trametinib (MEKi, Novartis, 2.5 mg/kg, oral gavage three times weekly), JAK/STAT3 inhibitor, ruxolitinib (STAT3i, TargetMol, 15 mg/kg, oral gavage three times weekly), αPD-1 antibody (BioXCell, Clone #BE0273, 200 µg/mouse, intraperitoneal injection twice weekly), or combined MEKi+STAT3i or MEKi+STAT3i with αPD1 beginning at 4-4.5 weeks of age. Mice were sacrificed after four weeks of treatment or when moribund. For survival studies, MEKi (2.5 mg/kg) and STAT3i (15 mg/kg) were administered by oral gavage 5 days/week starting at 4.5 weeks of age; αPD-1 antibody dosing remained intraperitoneal injection twice weekly. Six weeks after treatment initiation, the frequencies of MEKi+STAT3i and αPD1 antibody dosing were reduced to three times weekly and once weekly, respectively, and continued until mice were moribund (**Fig. S10).** T-cell depletion experiments are described in **Supplementary Methods**.

### Single cell RNA sequencing

The 10x Genomics Chromium Single Cell 3’ Reagent v3.1(Cat # PN-1000268) was used with standard conditions and volumes to process cell suspensions for 3’ transcriptional profiling. Single cell suspensions from PKT mice were extracted, and live cells were sorted using flow cytometry, and volumes were calculated for a target cell recovery of 100,000 cells and loaded on the Chromium Controller per manufacturer’s guidelines. The resultant purified cDNAs were quantified and qualitative assessed on the Agilent Bioanalyzer using the High Sensitivity DNA Kit (Cat #5067-4626). The final single cell 3’ libraries were quantified using the Qubit dsDNA High Sensitivity (Cat #Q33231) and qualitatively evaluated on the Agilent Bioanalyzer using the High Sensitivity DNA Kit . For sequencing, libraries were loaded at optimized concentrations onto an Illumina NovaSeq and paired-end sequenced under recommended settings (R1: 28 cycles; i7 index: 10 cycles; i5 index: 10 cycles; R2: 90 cycles). The libraries were diluted to varying nM concentrations in Illumina Resuspension Buffer (PN-15026770), denatured according to Illumina standard guidelines, and loaded on the Illumina NovaSeq at 1.2 nanomolar. The resulting intensity files were demultiplexed as FASTQ files using Illumina BaseSpace software and then aligned to the transcriptome using the 10x Genomics Cellranger (ver4.0.0) software package.

### Cluster identification and annotation in scRNAseq dataset

Principal component analysis was performed on the scaled data to reduce the dimensions, with number of components chosen based on a cumulative proportion (accumulated amount of explained variance) of 90%. Cells with >5% mitochondrial counts, less than 200 or more than 2500 unique feature counts were filtered out. Clusters of cells were identified by shared nearest neighbor algorithm. Cell type annotations were assigned to clusters based on the expression of canonical features in a minimum percentage of cells.

### Differential gene expression and pathway analysis in scRNAseq dataset

Prior to differential gene expression analysis, each cluster of interest was subjected to normalization, scaling, and PCA. Next, the function “FindMarkers” from Seurat v4.0 R package was utilized to finds the differentially expressed genes for identity classes (MEKi/STAT3i vs. vehicle). Genes were considered differentially expressed if detected in at least 25% of clusters, with default log fold change of 0.25. Wilcoxon Rank Sum test was used.

Gene set enrichment analysis (GSEA) for pathway analysis was performed using the “fgsea” R package on differentially expressed genes (log(FC) >0.5, and false discovery rate (FDR) adjusted p-value<0.05). “msigdbr” R package was utilized to access the following databases: C2 (KEGG, REACTOME, PID, BIOCARTA), and C5 (GO:BP). Among the different databases, the gene sets that met the statistical requirements were then curated, visualized via “ggplot2” R package, and ordered by Normalized Enrichment Score (NES).

### Pseudotime analysis

Pseudotime analysis was performed with “Monocle3” R package on scRNAseq data. CAF Seurat object was imported and transformed to CellDataSet (cds). CAF1 was chosen to act as the starting cluster, based on shared gene signature by Hosein *et al* (*26*). Single-cell trajectories was constructed, overlaid on a UMAP of the CAF cluster, and minor bifurcations mapped.

### CRISPR/Cas9 gene editing in KPC CAFs

To genetically ablate *Mek1* and *Stat3* in PDPN^+^ CAFs isolated from KPC mice (see **Supplementary Methods**), transEDIT^TM^ Lentiviral gRNA target gene sets specific for mouse *Map2k1* and *Stat3* (Cat. #CCMV 1101) were used (**Table S3**). CAFs were transduced with these lentiviral particles in conjugation with a Cas9 expressing vector (pCLIP-Cas9-nuclease vector). Cells were kept in appropriate antibiotic selection (puromycin or blasticidin) for the enrichment of transduced cells. To further increase genome editing efficiency, cells transduced with CRISPR/Cas9 vectors expressing a fluorescent protein were subjected to FACS to seed single cells into a 96 well plate at a density of 1-2.0 cells per well in 100 ul/ well of a complete culture medium. Adherent cells grown out from one colony (single clonal population) were identified using microscopy and expanded. Ensuing CAF-EV, CAF-*Mek1*^KO^, CAF-*Stat3*^KO^, CAF-*Mek1*^KO^*Stat3*^KO^ constructs were validated for knockout of respective genes by western blot analysis using antibodies specific for total MEK1 (CST#12671S) and STAT3 protein (CST#9139S) (**Fig. S4**).

### Bulk RNA sequencing and analysis of CRISPR/Cas9-edited CAF constructs

Purified RNA was obtained from *in vitro* CAF-EV, CAF-*Mek1*^KO^, CAF-*Stat3*^KO^, CAF-*Mek1*^KO^*Stat3*^KO^ cells in triplicates using RNeasy Kit (Qiagen) according to the manufacturer’s protocol. RNA quality was assessed on a bioanalyzer using the Agilent RNA 6000 Nano kit (5067-1511; Agilent Technologies); 0.2-1 μg of RNA with RNA integrity number > 7 proceeded for library preparation using an Illumina TruSeq RNA prep kit. Libraries were then sequenced using Illumina Nova Seq SP300. Reads from RNA-Seq were mapped to the Mus musculus genome (GRCm38) using STAR (ver.2.5.0) aligner. Raw counts were generated based on Ensembl genes (GENCODE M13) with feature Counts (ver.1.5.0). DE genes were identified using DESeq2 and determined by a threshold of FDR-corrected p-value<0.05. Pathway analysis was performed using Gene Set Enrichment Analysis with pathway database: Canonical Pathway (ver.5.1) provided by MSigDB database at Broad Institute was also used for the analysis.

### Flow cytometry

Pancreatic tumors harvested from PKT mice were enzymatically digested using solution of 0.6 mg/ml of collagenase P (Roche), 0.8 mg/ml Collagenase V (Sigma Aldrich), 0.6 mg/ml soybean trypsin inhibitor (Sigma Aldrich), and 1800 U/ml DNase I (Thermo Scientific) in RPMI medium for 20-30 minutes at 37°C. Samples were then washed and resuspended in cold PEB solution followed by straining through 40 µm mesh filter to obtain single cell suspension. Spleen samples were processed using RBC lysis buffer, followed passage through 100 µm mesh filter. Samples were frozen at -80°C until further use. Prior to flow cytometry staining, samples were thawed, washed prior to incubation with FcR blocking reagent (Miltenyi Biotec), and subsequently stained with fluorescently conjugated antibodies listed in **Table S1.** Live/dead cell discrimination was performed using Live/Dead Aqua (Life Technologies) or Ghost Red Dye 780 (TONBObiosciences) as per manufacturer’s protocol. Cell fixation was performed using 1% formaldehyde solution (Thermo Fisher). Flow cytometric data acquisition was performed on CytoFLEX S (Beckman Coulter) and analyzed using FlowJo v10 software (BD Life Sciences). Gating strategies are depicted in **Fig. S12**.

### Time of flight mass cytometry (CyTOF)

Single-cell suspensions from PKT tumors were flow sorted using FACS Aria III (BD Life Sciences) to remove non-cellular contaminants. Samples were resuspended and viability staining performed using 2.5 µM cisplatin (Fluidigm) working solution following by Fc receptor blocking and surface staining for 30 min at room temperature using tagged antibodies as shown in **Table S2**. Fixation was then performed using 1.6% formaldehyde solution (Thermo Fisher). Cells were suspended in 125 nM Cell-ID Intercalator-Ir solution (Fluidigm) overnight at 4°C. The following day, cells were washed and resuspended in 0.1X EQ Four Element Calibration Beads (Fluidigm) before acquisition. Data were acquired by Helios Mass Cytometer (Fluidigm) and analyzed using FlowJo v10 and Cytobank (Beckman Coulter). To generate viSNE plots, individual files from each treatment group were concatenated into a single file and then a subset of equal events were selected at random through FlowJo DownSample plugin. Total number of CD45^+^ cells from each group were used to create viSNE through Cytobank with 2000 iterations, perplexity of 30, and theta of 0.5.

### Histologic Analysis, Western Blotting, qPCR

Histologic analysis, western blotting, and qPCR analysis was performed as described previously (*22, 23, 51*). For details, refer to **Supplementary Methods**.

### T-cell Functional Cytotoxicity Assays

PKT mice were treated with vehicle, αPD-1, or MEKi+STAT3i with αPD-1 for 4 weeks before sacrifice and processing of tumor samples for flow cytometry. FACS-sorted viable EpCAM^+^ tumor cells and total splenocytes were isolated from each mouse. Tumor cells were irradiated (4 Gy for 7 minutes) and plated in a direct co-culture system with isolated splenocytes. Condition media was collected after 72 hours of co-culture and protein content determined by BCA method. Secreted IFN-γ was measured by ELISA as per manufacturer’s instruction (R&D Systems).

For measurement of intratumoral Granzyme B (GzB) levels, pancreatic tumor homogenates were prepared from PKT mice treated with vehicle, αPD-1, or MEKi+STAT3i with αPD-1 for 4 weeks. Tumor lysates were quantified and 40 μg protein loaded for analysis of GzB levels by ELISA (R&D Systems).

### Statistical analysis

Descriptive statistics were calculated using Prism 9.0 (GraphPad Software). Results are shown as mean±SEM. Multiple comparisons were performed using ANOVA followed by Tukey’s multiple comparisons test. The paired two-sided Student’s t test was used for two-group comparison. Survival curves were estimated using the Kaplan-Meier method and differences between groups were assessed using the log-rank test. An α-cutoff ≤0.05 was used to define statistical significance.

## SUPPLEMENTARY MATERIALS

Supplementary Methods

Figures S1 to S12 with adjoining legends

Table S1 to S6 with adjoining legends

## Author Contributions

Conception and design: JD, XD, AB, PL, NBM

Development of methodology: JD, XD, AB, IDS, PL, SM, ARD, VTG, FM, PJH, NSN, NBM

Acquisition of data (provided animals, acquired and managed patients, provided facilities, etc.): JD, XD, AB, IDS, PL, SM, ARD, SS, VTG, ZZ, FM, OU, YB, PJH, NSN, NBM

Analysis and interpretation of data (e.g., statistical analysis, biostatistics, computational analysis): JD, XD, AB, PL, SM, VTG, FM, OU, YB, NSN, NBM

Writing, review, and/or revision of the manuscript: JD, XD, AB, IDS, PL, SM, VTG, ARD, SS, ZZ, FM, OU, YB, PJH, NSN, NBM

Administrative, technical, or material support (i.e., reporting or organizing data, constructing databases): JD, AB, YB, NSN, NBM

Study supervision: JD, NBM

## Data and Materials Availability

Single cell RNA sequencing matrix files will be deposited in NIH GEO database upon acceptance of the manuscript. All other data are available in the main text or the supplementary materials.

## REFERENCES

1. M. A. Collins, F. Bednar, Y. Zhang, J. C. Brisset, S. Galban, C. J. Galban, S. Rakshit, K. S. Flannagan, N. V. Adsay, M. Pasca di Magliano, Oncogenic Kras is required for both the initiation and maintenance of pancreatic cancer in mice. J Clin Invest 122, 639–653 (2012).

2. E. M. Kerr, E. Gaude, F. K. Turrell, C. Frezza, C. P. Martins, Mutant Kras copy number defines metabolic reprogramming and therapeutic susceptibilities. Nature 531, 110–113 (2016).

3. G. L. Beatty, S. Eghbali, R. Kim, Deploying Immunotherapy in Pancreatic Cancer: Defining Mechanisms of Response and Resistance. Am Soc Clin Oncol Educ Book 37, 267–278 (2017).

4. J. Tang, J. X. Yu, V. M. Hubbard-Lucey, S. T. Neftelinov, J. P. Hodge, Y. Lin, Trial watch: The clinical trial landscape for PD1/PDL1 immune checkpoint inhibitors. Nat Rev Drug Discov 17, 854–855 (2018).

5. A. Marabelle, D. T. Le, P. A. Ascierto, A. M. Di Giacomo, A. De Jesus-Acosta, J. P. Delord, R. Geva, M. Gottfried, N. Penel, A. R. Hansen, S. A. Piha-Paul, T. Doi, B. Gao, H. C. Chung, J. Lopez-Martin, Y. J. Bang, R. S. Frommer, M. Shah, R. Ghori, A. K. Joe, S. K. Pruitt, L. A. Diaz, Jr., Efficacy of Pembrolizumab in Patients With Noncolorectal High Microsatellite Instability/Mismatch Repair-Deficient Cancer: Results From the Phase II KEYNOTE-158 Study. J Clin Oncol 38, 1–10 (2020).

6. E. M. O’Reilly, D. Y. Oh, N. Dhani, D. J. Renouf, M. A. Lee, W. Sun, G. Fisher, A. Hezel, S. C. Chang, G. Vlahovic, O. Takahashi, Y. Yang, D. Fitts, P. A. Philip, Durvalumab With or Without Tremelimumab for Patients With Metastatic Pancreatic Ductal Adenocarcinoma: A Phase 2 Randomized Clinical Trial. JAMA Oncol, (2019).

7. V. P. Balachandran, G. L. Beatty, S. K. Dougan, Broadening the Impact of Immunotherapy to Pancreatic Cancer: Challenges and Opportunities. Gastroenterology 156, 2056–2072 (2019).

8. P. P. Provenzano, C. Cuevas, A. E. Chang, V. K. Goel, D. D. Von Hoff, S. R. Hingorani, Enzymatic targeting of the stroma ablates physical barriers to treatment of pancreatic ductal adenocarcinoma. Cancer Cell 21, 418–429 (2012).

9. C. X. Dominguez, S. Muller, S. Keerthivasan, H. Koeppen, J. Hung, S. Gierke, B. Breart, O. Foreman, T. W. Bainbridge, A. Castiglioni, Y. Senbabaoglu, Z. Modrusan, Y. Liang, M. R. Junttila, C. Klijn, R. Bourgon, S. J. Turley, Single-Cell RNA Sequencing Reveals Stromal Evolution into LRRC15(+) Myofibroblasts as a Determinant of Patient Response to Cancer Immunotherapy. Cancer Discov 10, 232–253 (2020).

10. A. N. Hosein, R. A. Brekken, A. Maitra, Pancreatic cancer stroma: an update on therapeutic targeting strategies. Nat Rev Gastroenterol Hepatol 17, 487–505 (2020).

11. C. Feig, J. O. Jones, M. Kraman, R. J. Wells, A. Deonarine, D. S. Chan, C. M. Connell, E. W. Roberts, Q. Zhao, O. L. Caballero, S. A. Teichmann, T. Janowitz, D. I. Jodrell, D. A. Tuveson, D. T. Fearon, Targeting CXCL12 from FAP-expressing carcinoma-associated fibroblasts synergizes with anti-PD-L1 immunotherapy in pancreatic cancer. Proc Natl Acad Sci U S A 110, 20212–20217 (2013).

12. G. Biffi, T. E. Oni, B. Spielman, Y. Hao, E. Elyada, Y. Park, J. Preall, D. A. Tuveson, IL1-Induced JAK/STAT Signaling Is Antagonized by TGFbeta to Shape CAF Heterogeneity in Pancreatic Ductal Adenocarcinoma. Cancer Discov 9, 282–301 (2019).

13. A. R. Dosch, S. Singh, X. Dai, S. Mehra, I. C. Silva, A. Bianchi, S. Srinivasan, Z. Gao, Y. Ban, X. Chen, S. Banerjee, N. S. Nagathihalli, J. Datta, N. B. Merchant, Targeting Tumor-Stromal IL6/STAT3 Signaling through IL1 Receptor Inhibition in Pancreatic Cancer. Mol Cancer Ther 20, 2280–2290 (2021).

14. D. Ohlund, A. Handly-Santana, G. Biffi, E. Elyada, A. S. Almeida, M. Ponz-Sarvise, V. Corbo, T. E. Oni, S. A. Hearn, E. J. Lee, Chio, II, C. I. Hwang, H. Tiriac, L. A. Baker, D. D. Engle, C. Feig, A. Kultti, M. Egeblad, D. T. Fearon, J. M. Crawford, H. Clevers, Y. Park, D. A. Tuveson, Distinct populations of inflammatory fibroblasts and myofibroblasts in pancreatic cancer. J Exp Med 214, 579–596 (2017).

15. E. Elyada, M. Bolisetty, P. Laise, W. F. Flynn, E. T. Courtois, R. A. Burkhart, J. A. Teinor, P. Belleau, G. Biffi, M. S. Lucito, S. Sivajothi, T. D. Armstrong, D. D. Engle, K. H. Yu, Y. Hao, C. L. Wolfgang, Y. Park, J. Preall, E. M. Jaffee, A. Califano, P. Robson, D. A. Tuveson, Cross-Species Single-Cell Analysis of Pancreatic Ductal Adenocarcinoma Reveals Antigen-Presenting Cancer-Associated Fibroblasts. Cancer Discov 9, 1102–1123 (2019).

16. H. Huang, Y. Zhang, V. Gallegos, N. Sorrelle, M. M. Zaid, J. Toombs, W. Du, S. Wright, M. Hagopian, Z. Wang, A. N. Hosein, A. A. Sathe, C. Xing, E. J. Koay, K. E. Driscoll, R. A. Brekken, Targeting TGFbetaR2-mutant tumors exposes vulnerabilities to stromal TGFbeta blockade in pancreatic cancer. EMBO Mol Med 11, e10515 (2019).

17. Y. Shi, W. Gao, N. K. Lytle, P. Huang, X. Yuan, A. M. Dann, M. Ridinger-Saison, K. E. DelGiorno, C. E. Antal, G. Liang, A. R. Atkins, G. Erikson, H. Sun, J. Meisenhelder, E. Terenziani, G. Woo, L. Fang, T. P. Santisakultarm, U. Manor, R. Xu, C. R. Becerra, E. Borazanci, D. D. Von Hoff, P. M. Grandgenett, M. A. Hollingsworth, M. Leblanc, S. E. Umetsu, E. A. Collisson, M. Scadeng, A. M. Lowy, T. R. Donahue, T. Reya, M. Downes, R. M. Evans, G. M. Wahl, T. Pawson, R. Tian, T. Hunter, Targeting LIF-mediated paracrine interaction for pancreatic cancer therapy and monitoring. Nature 569, 131–135 (2019).

18. D. V. Catenacci, M. R. Junttila, T. Karrison, N. Bahary, M. N. Horiba, S. R. Nattam, R. Marsh, J. Wallace, M. Kozloff, L. Rajdev, D. Cohen, J. Wade, B. Sleckman, H. J. Lenz, P. Stiff, P. Kumar, P. Xu, L. Henderson, N. Takebe, R. Salgia, X. Wang, W. M. Stadler, F. J. de Sauvage, H. L. Kindler, Randomized Phase Ib/II Study of Gemcitabine Plus Placebo or Vismodegib, a Hedgehog Pathway Inhibitor, in Patients With Metastatic Pancreatic Cancer. J Clin Oncol 33, 4284–4292 (2015).

19. A. H. Ko, N. LoConte, M. A. Tempero, E. J. Walker, R. Kate Kelley, S. Lewis, W. C. Chang, E. Kantoff, M. W. Vannier, D. V. Catenacci, A. P. Venook, H. L. Kindler, A Phase I Study of FOLFIRINOX Plus IPI-926, a Hedgehog Pathway Inhibitor, for Advanced Pancreatic Adenocarcinoma. Pancreas 45, 370–375 (2016).

20. M. P. di Magliano, C. D. Logsdon, Roles for KRAS in pancreatic tumor development and progression. Gastroenterology 144, 1220–1229 (2013).

21. J. R. Infante, B. G. Somer, J. O. Park, C. P. Li, M. E. Scheulen, S. M. Kasubhai, D. Y. Oh, Y. Liu, S. Redhu, K. Steplewski, N. Le, A randomised, double-blind, placebo-controlled trial of trametinib, an oral MEK inhibitor, in combination with gemcitabine for patients with untreated metastatic adenocarcinoma of the pancreas. Eur J Cancer 50, 2072–2081 (2014).

22. N. S. Nagathihalli, J. Castellanos, P. Lamichhane, F. Messaggio, C. Shi, X. Dai, P. Rai, X. Chen, M. N. VanSaun, N. B. Merchant, Inverse Correlation of STAT3 and MEK Signaling Mediates Resistance to RAS Pathway Inhibition in Pancreatic Cancer. Cancer Res, (2018).

23. N. S. Nagathihalli, J. A. Castellanos, C. Shi, Y. Beesetty, M. L. Reyzer, R. Caprioli, X. Chen, A. J. Walsh, M. C. Skala, H. L. Moses, N. B. Merchant, Signal Transducer and Activator of Transcription 3, Mediated Remodeling of the Tumor Microenvironment Results in Enhanced Tumor Drug Delivery in a Mouse Model of Pancreatic Cancer. Gastroenterology 149, 1932–1943 e1939 (2015).

24. H. Ijichi, A. Chytil, A. E. Gorska, M. E. Aakre, Y. Fujitani, S. Fujitani, C. V. Wright, H. L. Moses, Aggressive pancreatic ductal adenocarcinoma in mice caused by pancreas-specific blockade of transforming growth factor-beta signaling in cooperation with active Kras expression. Genes Dev 20, 3147–3160 (2006).

25. V. Cremasco, J. L. Astarita, A. L. Grauel, S. Keerthivasan, K. MacIsaac, M. C. Woodruff, M. Wu, L. Spel, S. Santoro, Z. Amoozgar, T. Laszewski, S. C. Migoni, K. Knoblich, A. L. Fletcher, M. LaFleur, K. W. Wucherpfennig, E. Pure, G. Dranoff, M. C. Carroll, S. J. Turley, FAP Delineates Heterogeneous and Functionally Divergent Stromal Cells in Immune-Excluded Breast Tumors. Cancer Immunol Res 6, 1472–1485 (2018).

26. A. N. Hosein, H. Huang, Z. Wang, K. Parmar, W. Du, J. Huang, A. Maitra, E. Olson, U. Verma, R. A. Brekken, Cellular heterogeneity during mouse pancreatic ductal adenocarcinoma progression at single-cell resolution. JCI Insight 5, (2019).

27. T. Xie, Y. Wang, N. Deng, G. Huang, F. Taghavifar, Y. Geng, N. Liu, V. Kulur, C. Yao, P. Chen, Z. Liu, B. Stripp, J. Tang, J. Liang, P. W. Noble, D. Jiang, Single-Cell Deconvolution of Fibroblast Heterogeneity in Mouse Pulmonary Fibrosis. Cell Rep 22, 3625–3640 (2018).

28. Y. Mizutani, H. Kobayashi, T. Iida, N. Asai, A. Masamune, A. Hara, N. Esaki, K. Ushida, S. Mii, Y. Shiraki, K. Ando, L. Weng, S. Ishihara, S. M. Ponik, M. W. Conklin, H. Haga, A. Nagasaka, T. Miyata, M. Matsuyama, T. Kobayashi, T. Fujii, S. Yamada, J. Yamaguchi, T. Wang, S. L. Woods, D. Worthley, T. Shimamura, M. Fujishiro, Y. Hirooka, A. Enomoto, M. Takahashi, Meflin-Positive Cancer-Associated Fibroblasts Inhibit Pancreatic Carcinogenesis. Cancer Res 79, 5367–5381 (2019).

29. X. Qiu, A. Hill, J. Packer, D. Lin, Y. A. Ma, C. Trapnell, Single-cell mRNA quantification and differential analysis with Census. Nature methods 14, 309–315 (2017).

30. T. D. Somerville, G. Biffi, J. Dassler-Plenker, S. K. Hur, X. Y. He, K. E. Vance, K. Miyabayashi, Y. Xu, D. Maia-Silva, O. Klingbeil, O. E. Demerdash, J. B. Preall, M. A. Hollingsworth, M. Egeblad, D. A. Tuveson, C. R. Vakoc, Squamous trans-differentiation of pancreatic cancer cells promotes stromal inflammation. Elife 9, (2020).

31. M. Buxade, H. Huerga Encabo, M. Riera-Borrull, L. Quintana-Gallardo, P. Lopez-Cotarelo, M. Tellechea, S. Martinez-Martinez, J. M. Redondo, J. Martin-Caballero, J. M. Flores, E. Bosch, J. L. Rodriguez-Fernandez, J. Aramburu, C. Lopez-Rodriguez, Macrophage-specific MHCII expression is regulated by a remote Ciita enhancer controlled by NFAT5. J Exp Med 215, 2901–2918 (2018).

32. L. Li, X. Guo, X. Shi, C. Li, W. Wu, C. Yan, H. Wang, H. Li, C. Xu, Ionic CD3-Lck interaction regulates the initiation of T-cell receptor signaling. Proc Natl Acad Sci U S A 114, E5891–E5899 (2017).

33. N. Ryan, K. Anderson, G. Volpedo, O. Hamza, S. Varikuti, A. R. Satoskar, S. Oghumu, STAT1 inhibits T-cell exhaustion and myeloid derived suppressor cell accumulation to promote antitumor immune responses in head and neck squamous cell carcinoma. Int J Cancer 146, 1717–1729 (2020).

34. J. Bae, A. Munshi, C. Li, M. Samur, R. Prabhala, C. Mitsiades, K. C. Anderson, N. C. Munshi, Heat shock protein 90 is critical for regulation of phenotype and functional activity of human T lymphocytes and NK cells. J Immunol 190, 1360–1371 (2013).

35. R. Winograd, K. T. Byrne, R. A. Evans, P. M. Odorizzi, A. R. Meyer, D. L. Bajor, C. Clendenin, B. Z. Stanger, E. E. Furth, E. J. Wherry, R. H. Vonderheide, Induction of T-cell Immunity Overcomes Complete Resistance to PD-1 and CTLA-4 Blockade and Improves Survival in Pancreatic Carcinoma. Cancer Immunol Res 3, 399–411 (2015).

36. A. Azad, S. Yin Lim, Z. D’Costa, K. Jones, A. Diana, O. J. Sansom, P. Kruger, S. Liu, W. G. McKenna, O. Dushek, R. J. Muschel, E. Fokas, PD-L1 blockade enhances response of pancreatic ductal adenocarcinoma to radiotherapy. EMBO Mol Med 9, 167–180 (2017).

37. H. Jiang, S. Hegde, B. L. Knolhoff, Y. Zhu, J. M. Herndon, M. A. Meyer, T. M. Nywening, W. G. Hawkins, I. M. Shapiro, D. T. Weaver, J. A. Pachter, A. Wang-Gillam, D. G. DeNardo, Targeting focal adhesion kinase renders pancreatic cancers responsive to checkpoint immunotherapy. Nat Med 22, 851–860 (2016).

38. K. C. Soares, A. A. Rucki, A. A. Wu, K. Olino, Q. Xiao, Y. Chai, A. Wamwea, E. Bigelow, E. Lutz, L. Liu, S. Yao, R. A. Anders, D. Laheru, C. L. Wolfgang, B. H. Edil, R. D. Schulick, E. M. Jaffee, L. Zheng, PD-1/PD-L1 blockade together with vaccine therapy facilitates effector T-cell infiltration into pancreatic tumors. J Immunother 38, 1–11 (2015).

39. C. W. Steele, S. A. Karim, J. D. G. Leach, P. Bailey, R. Upstill-Goddard, L. Rishi, M. Foth, S. Bryson, K. McDaid, Z. Wilson, C. Eberlein, J. B. Candido, M. Clarke, C. Nixon, J. Connelly, N. Jamieson, C. R. Carter, F. Balkwill, D. K. Chang, T. R. J. Evans, D. Strathdee, A. V. Biankin, R. J. B. Nibbs, S. T. Barry, O. J. Sansom, J. P. Morton, CXCR2 Inhibition Profoundly Suppresses Metastases and Augments Immunotherapy in Pancreatic Ductal Adenocarcinoma. Cancer Cell 29, 832–845 (2016).

40. L. Strauss, M. A. A. Mahmoud, J. D. Weaver, N. M. Tijaro-Ovalle, A. Christofides, Q. Wang, R. Pal, M. Yuan, J. Asara, N. Patsoukis, V. A. Boussiotis, Targeted deletion of PD-1 in myeloid cells induces antitumor immunity. Sci Immunol 5, (2020).

41. M. Ligorio, S. Sil, J. Malagon-Lopez, L. T. Nieman, S. Misale, M. Di Pilato, R. Y. Ebright, M. N. Karabacak, A. S. Kulkarni, A. Liu, N. Vincent Jordan, J. W. Franses, J. Philipp, J. Kreuzer, N. Desai, K. S. Arora, M. Rajurkar, E. Horwitz, A. Neyaz, E. Tai, N. K. C. Magnus, K. D. Vo, C. N. Yashaswini, F. Marangoni, M. Boukhali, J. P. Fatherree, L. J. Damon, K. Xega, R. Desai, M. Choz, F. Bersani, A. Langenbucher, V. Thapar, R. Morris, U. F. Wellner, O. Schilling, M. S. Lawrence, A. S. Liss, M. N. Rivera, V. Deshpande, C. H. Benes, S. Maheswaran, D. A. Haber, C. Fernandez-Del-Castillo, C. R. Ferrone, W. Haas, M. J. Aryee, D. T. Ting, Stromal Microenvironment Shapes the Intratumoral Architecture of Pancreatic Cancer. Cell 178, 160–175 e127 (2019).

42. W. Hugo, J. M. Zaretsky, L. Sun, C. Song, B. H. Moreno, S. Hu-Lieskovan, B. Berent-Maoz, J. Pang, B. Chmielowski, G. Cherry, E. Seja, S. Lomeli, X. Kong, M. C. Kelley, J. A. Sosman, D. B. Johnson, A. Ribas, R. S. Lo, Genomic and Transcriptomic Features of Response to Anti-PD-1 Therapy in Metastatic Melanoma. Cell 168, 542 (2017).

43. N. Riaz, J. J. Havel, V. Makarov, A. Desrichard, W. J. Urba, J. S. Sims, F. S. Hodi, S. Martin-Algarra, R. Mandal, W. H. Sharfman, S. Bhatia, W. J. Hwu, T. F. Gajewski, C. L. Slingluff, Jr., D. Chowell, S. M. Kendall, H. Chang, R. Shah, F. Kuo, L. G. T. Morris, J. W. Sidhom, J. P. Schneck, C. E. Horak, N. Weinhold, T. A. Chan, Tumor and Microenvironment Evolution during Immunotherapy with Nivolumab. Cell 171, 934–949 e916 (2017).

44. Y. Kieffer, H. R. Hocine, G. Gentric, F. Pelon, C. Bernard, B. Bourachot, S. Lameiras, L. Albergante, C. Bonneau, A. Guyard, K. Tarte, A. Zinovyev, S. Baulande, G. Zalcman, A. Vincent-Salomon, F. Mechta-Grigoriou, Single-Cell Analysis Reveals Fibroblast Clusters Linked to Immunotherapy Resistance in Cancer. Cancer Discov 10, 1330–1351 (2020).

45. G. Sheng, The developmental basis of mesenchymal stem/stromal cells (MSCs). BMC Dev Biol 15, 44 (2015).

46. M. Kortylewski, M. Kujawski, T. Wang, S. Wei, S. Zhang, S. Pilon-Thomas, G. Niu, H. Kay, J. Mule, W. G. Kerr, R. Jove, D. Pardoll, H. Yu, Inhibiting Stat3 signaling in the hematopoietic system elicits multicomponent antitumor immunity. Nat Med 11, 1314–1321 (2005).

47. M. J. Allegrezza, M. R. Rutkowski, T. L. Stephen, N. Svoronos, A. Perales-Puchalt, J. M. Nguyen, K. K. Payne, S. Singhal, E. B. Eruslanov, J. Tchou, J. R. Conejo-Garcia, Trametinib Drives T-cell-Dependent Control of KRAS-Mutated Tumors by Inhibiting Pathological Myelopoiesis. Cancer Res 76, 6253–6265 (2016).

48. E. Van Cutsem, M. A. Tempero, D. Sigal, D. Y. Oh, N. Fazio, T. Macarulla, E. Hitre, P. Hammel, A. E. Hendifar, S. E. Bates, C. P. Li, S. R. Hingorani, C. de la Fouchardiere, A. Kasi, V. Heinemann, A. Maraveyas, N. Bahary, L. Layos, V. Sahai, L. Zheng, J. Lacy, J. O. Park, F. Portales, P. Oberstein, W. Wu, D. Chondros, A. J. Bullock, H. Investigators, Randomized Phase III Trial of Pegvorhyaluronidase Alfa With Nab-Paclitaxel Plus Gemcitabine for Patients With Hyaluronan-High Metastatic Pancreatic Adenocarcinoma. J Clin Oncol 38, 3185–3194 (2020).

49. J. L. Carstens, P. Correa de Sampaio, D. Yang, S. Barua, H. Wang, A. Rao, J. P. Allison, V. S. LeBleu, R. Kalluri, Spatial computation of intratumoral T cells correlates with survival of patients with pancreatic cancer. Nat Commun 8, 15095 (2017).

50. A. Chytil, M. A. Magnuson, C. V. Wright, H. L. Moses, Conditional inactivation of the TGF-beta type II receptor using Cre:Lox. Genesis 32, 73–75 (2002).

51. N. S. Nagathihalli, J. A. Castellanos, M. N. VanSaun, X. Dai, M. Ambrose, Q. Guo, Y. Xiong, N. B. Merchant, Pancreatic stellate cell secreted IL-6 stimulates STAT3 dependent invasiveness of pancreatic intraepithelial neoplasia and cancer cells. Oncotarget 7, 65982–65992 (2016).

