## Supplementary Materials for "Combined MEK and STAT3 inhibition reprograms stromal inflammation to overcome immunotherapy resistance in pancreatic cancer"

### SUPPLEMENTARY METHODS

#### *Fluorescence-activated cell sorting for single-cell RNA sequencing*

Pancreas tissue harvested from both vehicle- and MEKi+STAT3i-treated PKT mice were mechanically dissociated to generate single cell suspensions as described previously. Cells were then resuspended in 10 mLs of buffer (autoMACS® Rinsing Solution in 5% MACS® BSA) and incubated for 10 minutes at 4 °C with 4′,6-Diamidino-2-phenylindole dihydrochloride (Roche Diagnostics) 1:1000 and DRAQ5™ (Biolegend) 1:500 to stain cells for viability. Lived cells sorted by BD FACSAria™ Fusion were then resuspended in PBS (BioWhittaker, Inc) with 3% BSA (5% MACS® BSA) for single-cell RNA sequencing.

#### *Isolation and Characterization of LSL-Kras$^{G12D/+}$; Trp53$^{R172H/+}$; Pdx1$^{Cre/+}$ (KPC) Mouse derived CAFs*

Mouse CAFs were isolated from KPC mice with histologically proven PDAC. CAFs were isolated from primary tumors using a combination of outgrowth and clonal isolation. The edge of the tumor mass was minced and mechanically disaggregated in RPMI containing 1% FBS, 0.125 mg/ml collagenase (Sigma-Aldrich), and 0.125 mg/ml dispase (Life Technologies) for 0.5 h at 37°C in a thermomixer. The sample was trypsinized for 10 min then quenched in 10% FBS/RPMI. The pellet containing tumor pieces was plated on a 6-cm dish, and fibroblasts were allowed to grow out and attach to the plastic. To avoid cancer cell contamination, after the cell culture was established and passaged couple of times, cells were subjected to FACS using antibodies against fibroblast surface marker PDPN and tumor epithelial cell marker EpCAM. Flow sorted PDPN$^{+}$ and EpCAM$^{-}$ cell population was further expanded for subsequent downstream experiment analysis.

***T cell depletion experiments in PKT mice***

Starting at 4.5 weeks of age, tumor bearing PKT mice were treated with 200μg each of $CD4^+$ and $CD8^+$ T cell depleting antibodies (BioXCell, anti-CD4 Ab Clone GK1.5 and anti-CD8 Ab Clone 2.43) via intraperitoneal injection once daily on day 3 and 7 then once weekly thereafter. MEKi (2.5 mg/kg/day) and STAT3i (15 mg/kg/day) treatment was administered by oral gavage five times weekly for the first two weeks, then three times a week until mice were moribund **(Figure S8)**. At sacrifice, tumor weight was recorded. Tumor and splenic samples were processed for flow cytometry to confirm depletion of $CD4^+$ and $CD8^+$ populations.

***Histologic Analysis***

Pancreatic tumor tissues were fixed in 10% neutral-buffered formalin, embedded in paraffin, and sectioned. Hematoxylin and eosin (H&E), Sirius Red, Masson's Trichrome, Alcian Blue were performed as previously described (*22, 23, 51*). For immunohistochemistry (IHC) and immunofluorescence (IF) staining, tissue sections were mounted on glass slides and deparaffinized in xylene followed by rehydration using alcohol gradient. Antigen retrieval was performed by incubating samples in citrate buffer (0.01 M, pH 6:0) and heating. Sections were blocked using BlockAid™ (Thermo Fischer) to preclude non-specific binding. For IHC experiments, endogenous peroxidase activity was quenched by incubating with 3% $H_2O_2$ for ten minutes. Sections were then incubated with the primary antibodies listed in **Table S4** in a humidified chamber at 4℃ overnight. On the following day, slides were washed and developed using VECTASTAIN R Elite ABC HRP based kit (Vector) as per the manufacturer's protocol with diaminobenzidine (DAB) as the chromogen. Tissue sections were counterstained with Mayer's hematoxylin, mounted, and imaged using DM750 Leica microscope (Leica Microsystems). For IF experiments, bound primary antibody was detected by incubating species-specific Alexa Fluor 594 and/or Alexa Fluor 488 secondary antibodies (Thermo Fisher) at room temperature for 1 hour followed by nuclear counterstaining with Hoechst 33342 (Thermo Fisher). IF images were acquired using Leica DMi8 microscope system (Leica Microsystems). Quantitative histological analysis was performed by sampling multiple random, non-overlapping fields in each tissue section and quantified using ImageJ software (NIH) to measure the percent positive area. Area of normal pancreas relative to tumor were analyzed in PKT sections by manual quantification (ImageJ) of randomly selected fields.

#### *Western Blotting*

Pancreatic tissue samples were harvested from PKT mice and snap frozen in liquid nitrogen. Tumor samples were thawed and homogenized in RIPA lysis buffer (Cell Signaling) containing protease inhibitor cocktail (Millipore Sigma) and PhosSTOP phosphatase inhibitor (Millipore Sigma) prior to sonication. Homogenate was then centrifuged at 4°C to remove the cellular debris and protein concentration determined using BCA. Tumor lysates containing 30-40 µg of equal protein were separated using 4-20 % SDS PAGE Mini-PROTEAN TGX Stain-Free Gel (Bio-Rad) and transferred on Trans-Blot® Turbo™ Midi PVDF Transfer Packs (Bio-Rad) using Trans-Blot Turbo Transfer System (Bio-Rad). For immunodetection, membranes were incubated with primary antibodies listed in **Table S5** at 4°C overnight. Membranes were then washed and incubated with corresponding specific secondary antibodies conjugated with horseradish peroxidase (Jackson Immuno Research Laboratory). Immunoreactive bands were developed using Pierce ECL Western Blotting Substrate (Thermo Scientific) or SuperSignal™ West Pico PLUS Chemiluminescent Substrate (Thermo Scientific). Densitometric analysis for quantification of protein expression was performed using ImageJ software.

#### *Quantitative Polymerase Chain Reaction (qPCR)*

RNA was obtained from tumor samples using RNeasy Kit (Qiagen) according to manufacturer's protocol. RNA concentration and quality were determined using NanoDrop spectrophotometer 2000 (Thermo Fisher). 1.5-2 µg of RNA was reverse transcribed to cDNA using High-Capacity cDNA Reverse Transcription Kit with RNase Inhibitor (Applied Biosystems). qPCR was performed with gene-specific predesigned primers ($RT^2$ qPCR Primer Assay, Qiagen) and iQ™ SYBR® Green Supermix (BioRad). Gene expression was normalized to the housekeeping gene *18s* using ΔΔCT method and reported as fold change relative to control.

***Cytokine Array***

Conditioned media (CM) was collected from KPC CAFs (Wild type and $MS^{KO}$) after incubation in serum-free media (SFM) for 48 hours. CM collected was subjected to protein concentration measurement. An equal amount of protein (300 µg) was added for analysis by cytokine array analysis (ARY006, R&D Systems) as per the manufacturer's protocol.

### SUPPLEMENTARY FIGURES

Supplementary Fig S1

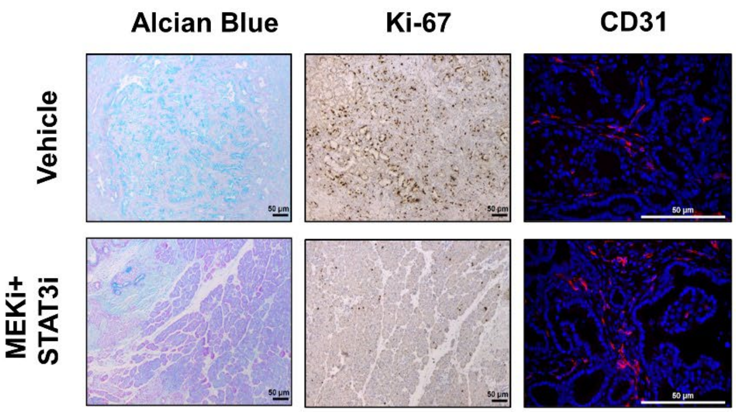


**Figure S1. Combined MEKi/STAT3i inhibition remodels PDAC stromal architecture.** Histologic analysis of pancreatic tumor tissue sections for Alcian blue, Ki67, and CD31 positive staining in PKT mice treated with vehicle and MEKi+STAT3i+αPD-1 for 4 weeks (n=3-5 mice/arm).

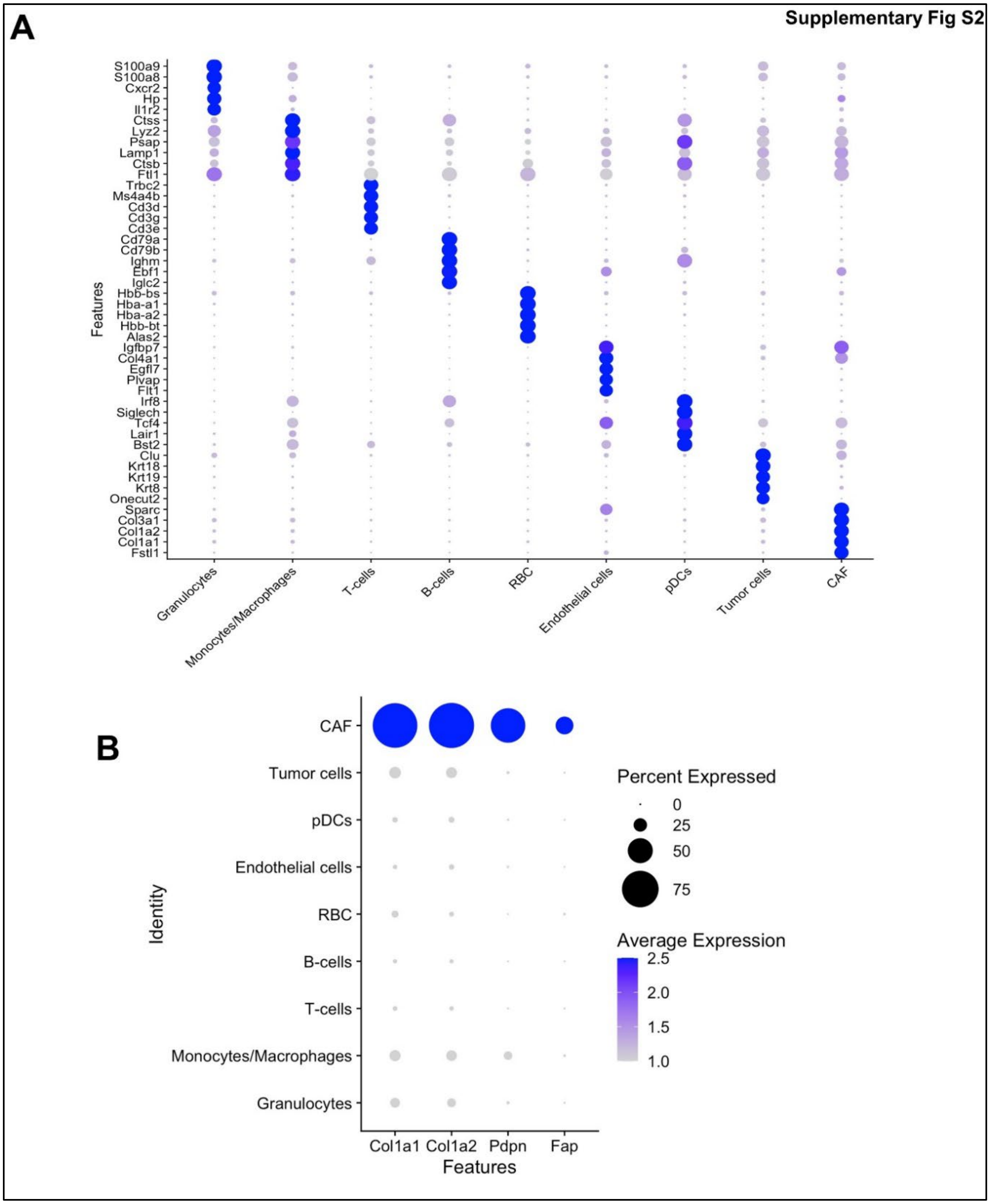


**Figure S2. Cellular identities identified in the PDAC TME via single-cell RNA sequencing in PKT mice. (A)** Bubble plot outlining the expression of canonical lineage cell markers utilized for cell cluster annotation in PKT scRNAseq dataset; **(B)** Canonical transcriptional descriptors of fibroblast identity *Col1a1, Col1a2, Fap, Pdpn* nominate the CAF sub-cluster in the scRNAseq dataset.

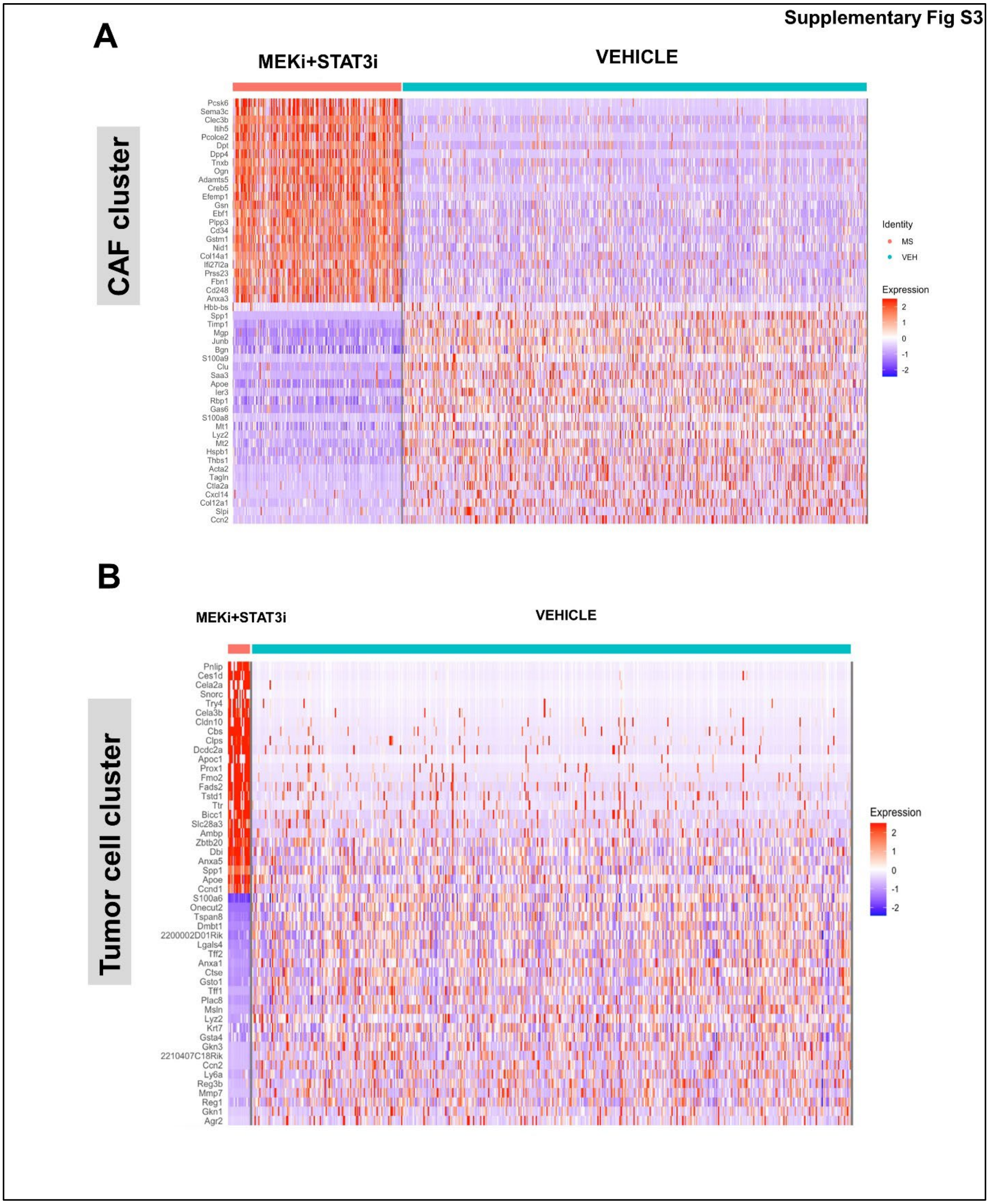


**Figure S3. Single cell RNA sequencing reveals reprogramming of the transcriptional machinery in tumor-cell and CAF compartments *in vivo*.** Top 25 differentially regulated genes from MEKi+STAT3i-treated vs. vehicle-treated PKT single-cell transcriptomes in: **(A)** Cancer-associated fibroblast; and **(B)** Tumor-cell subcluster.

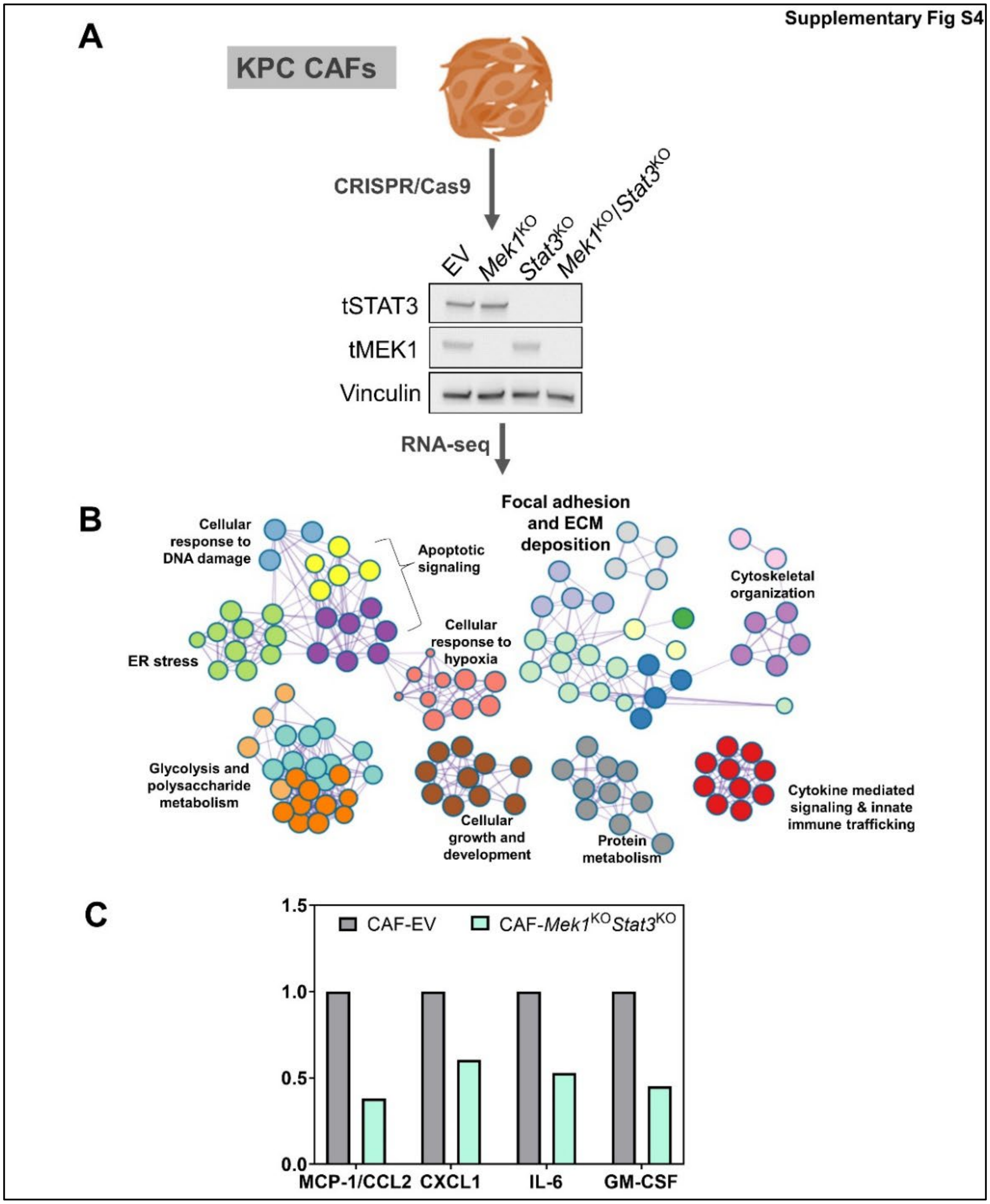


**Figure S4. CRISPR/cas9-enabled CAF-specific *Mek1* and *Stat3* genetic silencing modulates fibroinflammatory properties. (A)** Flow cytometry-sorted $FAP^+PDPN^+$ CAFs from 6 month-old $Kras^{G12D/+}$;$Trp53^{fl/+}$;$Pdx\text{-}^{Cre}$ (KPC) genetic mouse were subjected to CRISPR/Cas9 genetic editing with empty vector (EV) sgRNA, *Mek1* sgRNA alone, *Stat3* sgRNA alone, and combined *Mek1/Stat3* sgRNA (see Supplementary Methods for details). Target inhibition is shown via representative western blot; **(B)** Bulk RNA sequencing of all 4 CAF subsets was performed and differential gene set expression analysis of differentially regulated genes specific to the CAF-$Mek1^{KO}$ $Stat3^{KO}$ subset was subjected to Metascape network analysis to reveal several biologic processes. Differential gene ontologies are nominated by separate colors and labeled according to putative function; **(C)** CAF-EV and CAF-*Mek1*$^{KO}$ *Stat3*$^{KO}$ cells were expanded *in vitro* and conditioned media from each subjected to multiplex cytokine array. The histogram shows relative secretion of cytokines/chemokines implicated in myeloid cell chemoattraction in CAF-EV and CAF-*Mek1*$^{KO}$ *Stat3*$^{KO}$ cells.

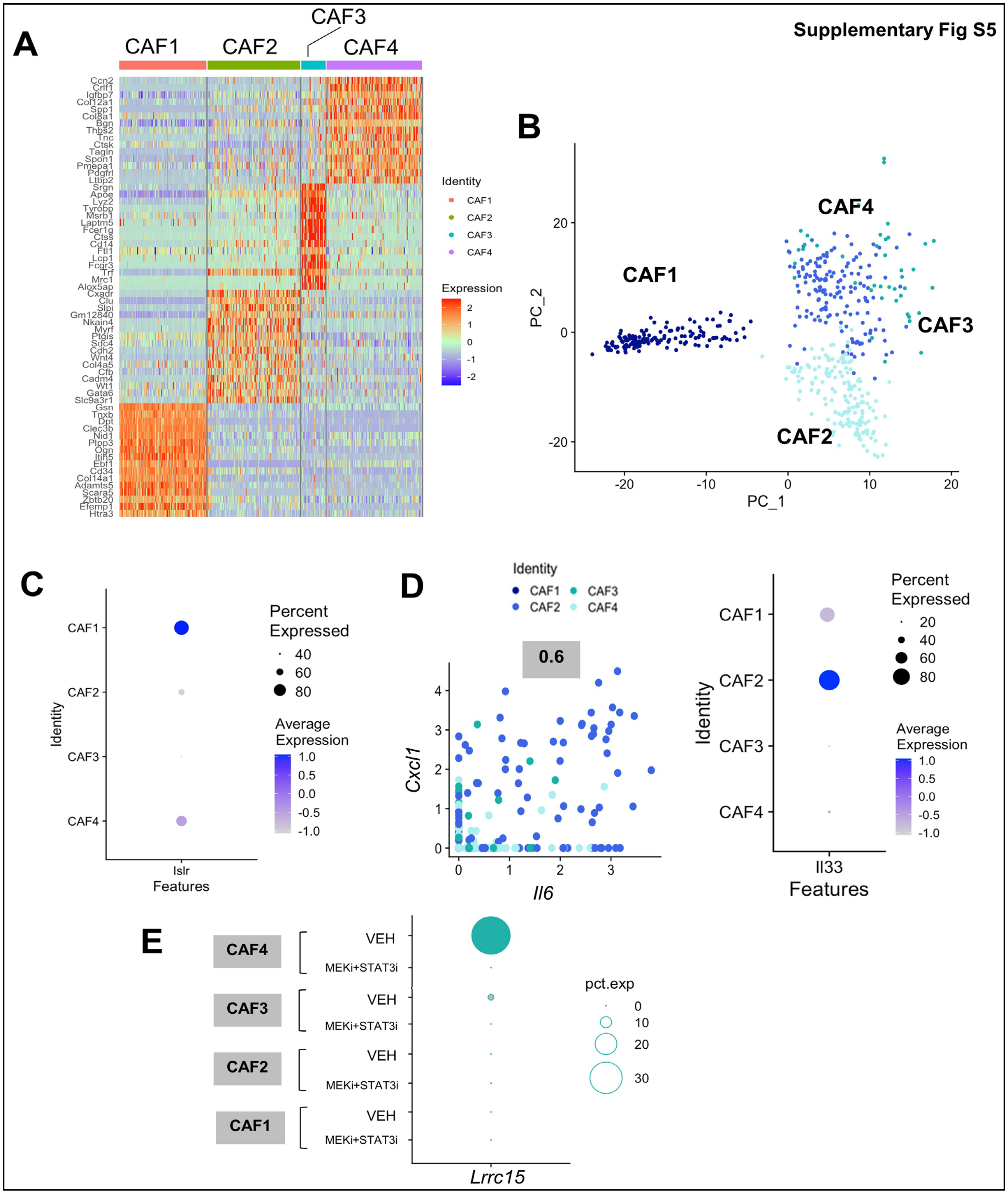


**Figure S5: Heterogeneity in transcriptional programs by single-cell RNA sequencing nominates four distinct CAF subsets in the PDAC TME. (A)** Heatmap showing top 15 differentially overexpressed genes distinguishing CAF1 though CAF4 sub-clusters in the scRNAseq dataset from PKT tumors. CAF sub-cluster identity is shown in color-coded header;

**(B)** Principal component analysis based on the most highly variable genes in the CAF scRNAseq cluster, as measured by variance-stabilizing transformation, depicts transcriptional divergence of CAF1 from CAF2-4 subsets; **(C)** Bubble plot showing relative expression of *Lslr* (Meflin)—a mesenchymal stromal/stem cell (MSC) marker recently implicated in tumor-restraining function in PDAC—across CAF1-4 subclusters; **(D)** Co-expression plot showing strength of co-expression of *Il6* and *Cxcl1* across CAF1-4 sub-cluster. Co-expression index of 0.6 is highest for CAF3 sub-cluster (*left*). Bubble plot shows relative expression of *Il33* (*right*), supporting its nomination as secretory/inflammatory CAF; **(E)** Dot plot depicting the expression of myCAF gene *Lrrc15* across different CAF sub-subsets comparing vehicle- and MEKi+STAT3i-treated cohorts.

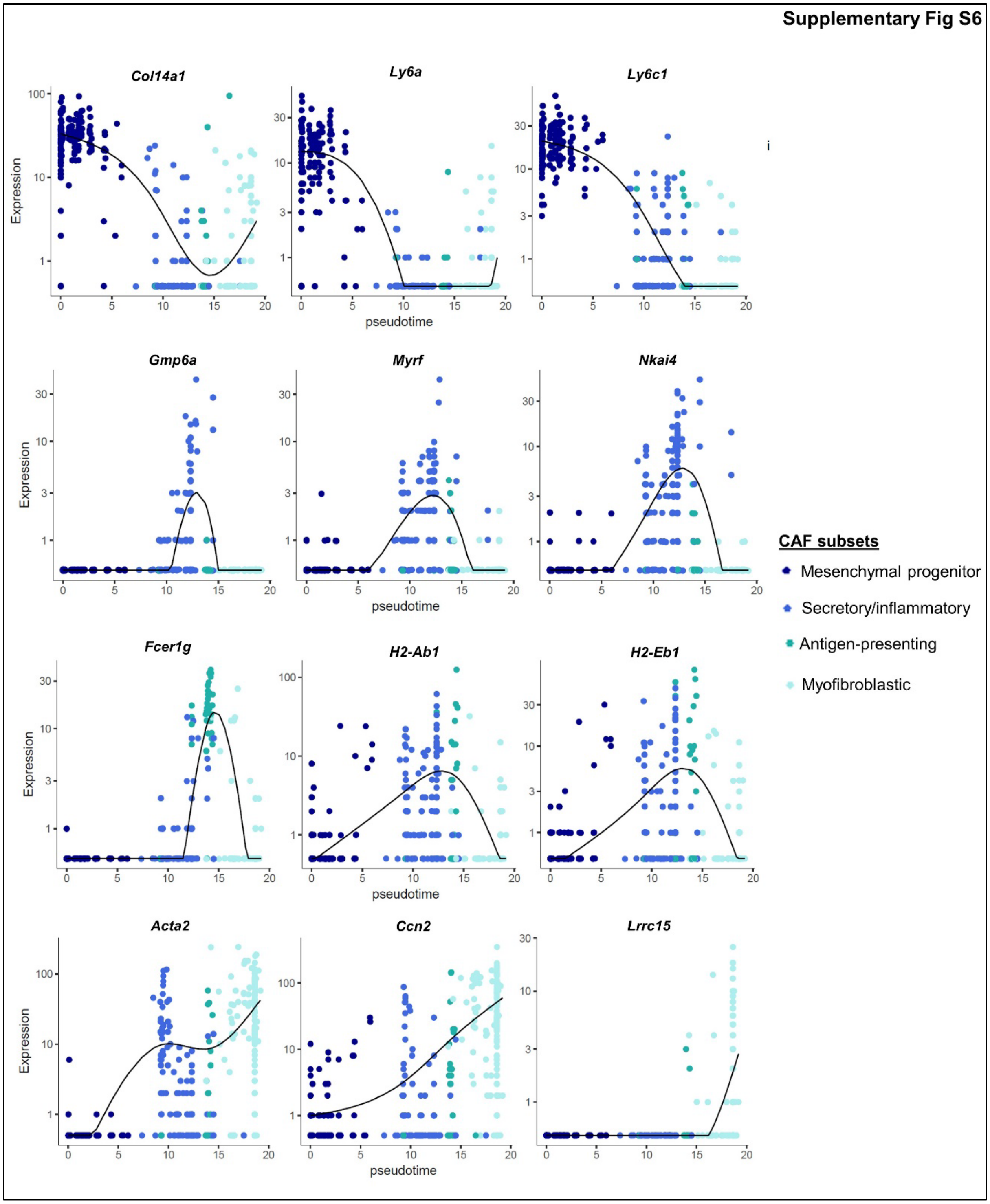


**Figure S6: Single-cell pseudotime lineage reconstruction reveals putative cancer-associated fibroblast developmental trajectories.** Gene expression dynamics of specific transcripts overexpressed in pseudotime reconstruction are shown across four putative CAF developmental states: mesenchymal progenitor (*Col14a1*, *Ly6a*, *Ly6c1*), secretory/inflammatory (*Gmp6a*, *Myrf*, *Nkai4*), antigen-presenting (*Fcer1g*, *H2-Ab1*, *H2-Eb1*), and myofibroblastic

(*Acta2*, *Ccn2*, *Lrrc15*). Relative expression density across pseudotimes is designated by solid black line.

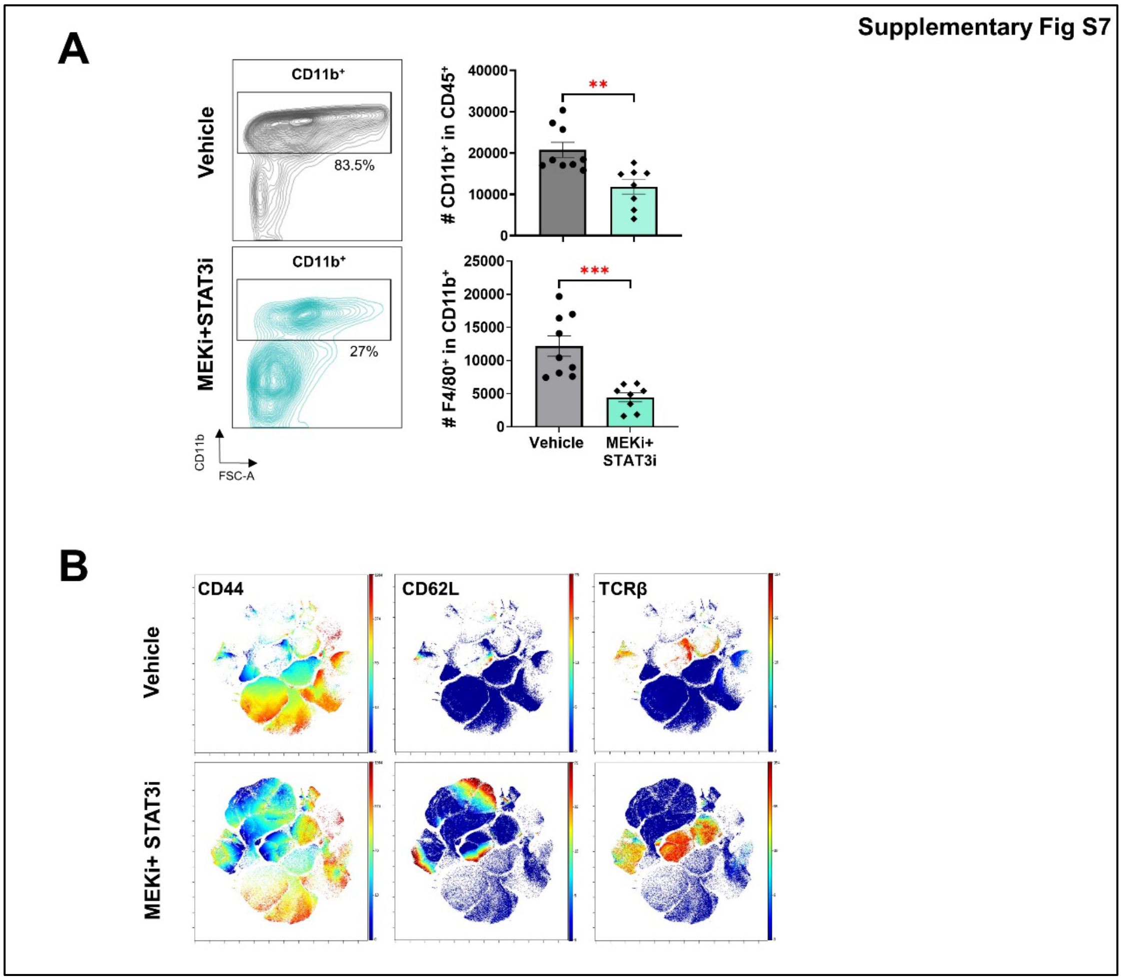


**Figure S7. Immune remodeling following MEKi+STAT3i in PKT mice. (A)** Flow cytometric profiling and quantification of absolute number of CD45⁺CD11b⁺ myeloid cells (representative contour plot shown on *left*) and CD11b⁺F4/80⁺ macrophages in MEKi+STAT3i-treated PKT tumors compared with vehicle-treated tumors; **(B)** viSNE plots of selected T-cell markers CD44, CD62L, and TCRβ from PKT tumors in mice treated with either vehicle or MEKi+STAT3i and immunophenotyped with CyTOF profiling. Events depicted are gated on viable CD45⁺ cells and show equal acquisition events. Scale bar = 50 μm. Data are shown as mean ± SEM. **, p<0.01; ***, p<0.001.

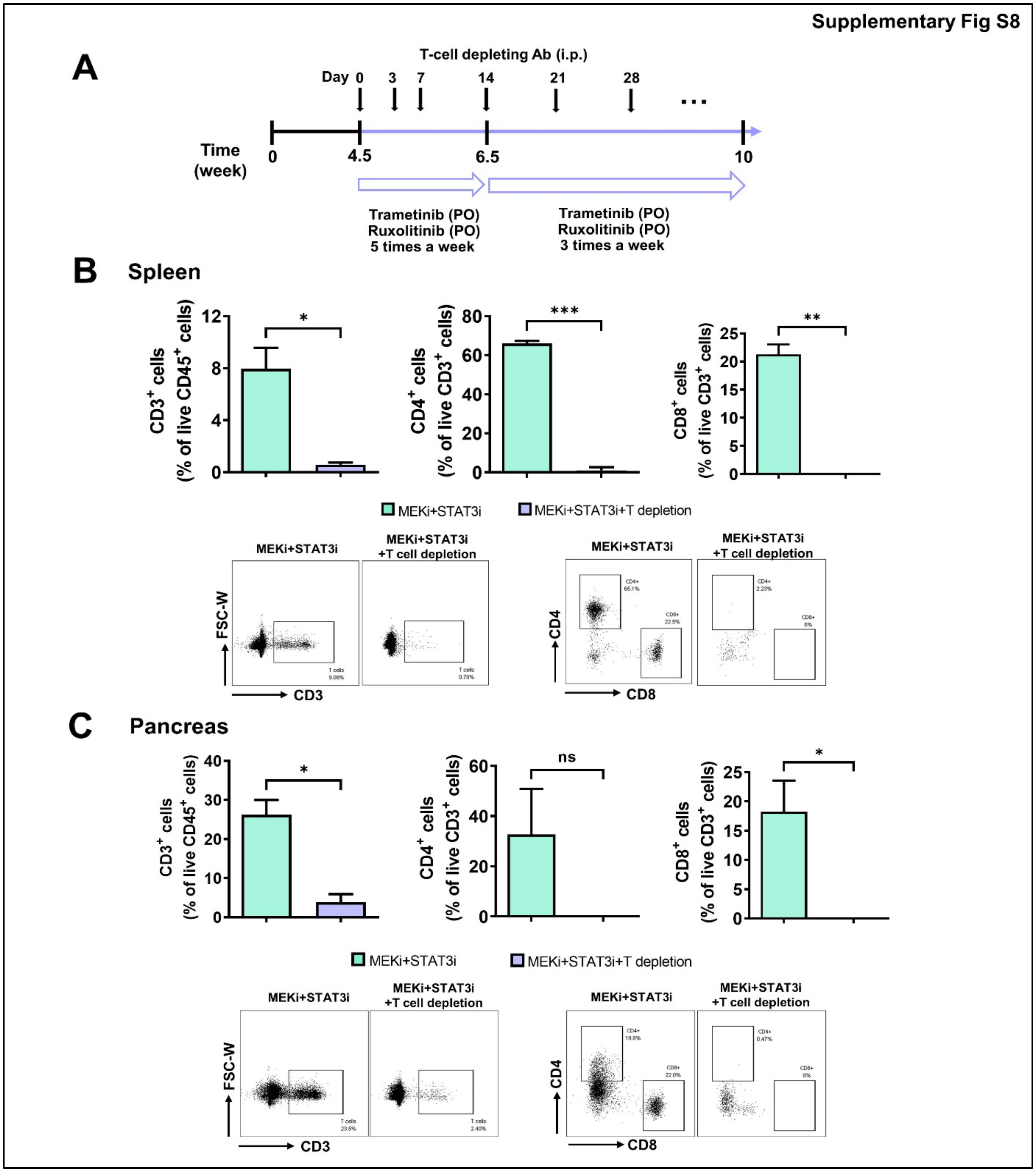


**Figure S8. T-cell depletion experiments in PKT mice. (A)** Treatment timeline for T-cell depletion experiments in PKT mice. Doses of Trametinib and Ruxolitinib are as defined in *Methods*. Flow cytometric analysis of **(B)** spleens and **(C)** pancreata isolated from PKT mice following treatment with MEKi/STAT3i as well as anti-CD4 and anti-CD8 antibodies demonstrating depletion of viable $CD3^+$, $CD4^+$, and $CD8^+$ cells compared to PKT mice treated with MEKi/STAT3i alone. Data are shown as mean ± SEM. ns, not significant; *, $p<0.05$; **, $p<0.01$; ***, $p<0.001$.

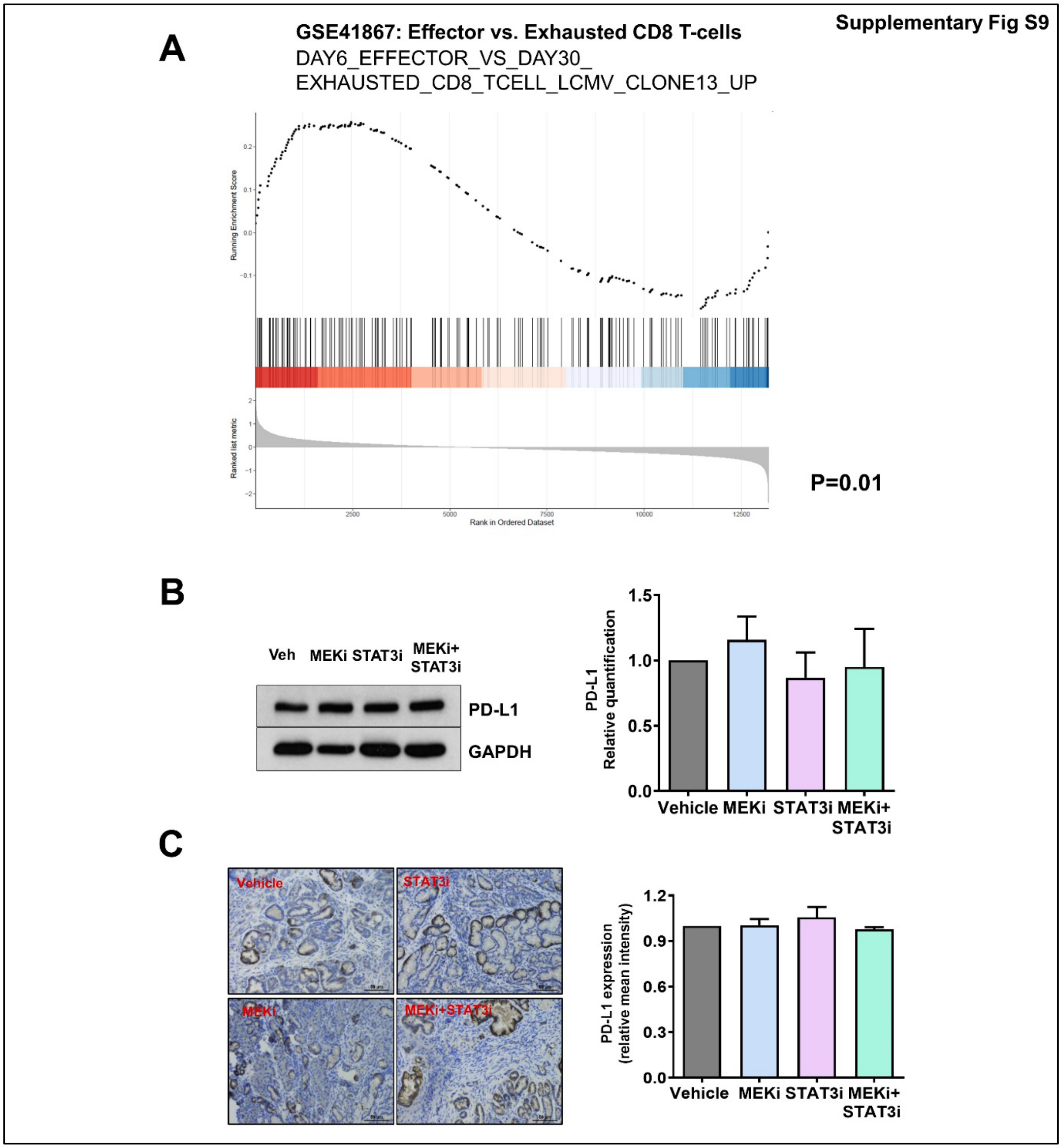


**Figure S9. T-cell activation and PD-L1 expression in the tumor microenvironment following MEKi+STAT3i treatment. (A)** Enrichment plot showing relative expression of genes in the GSE41867 pathway—denoting genes upregulated in effector compared with exhausted T-cells in chronic viral inflammation (LCMV) models—in MEKi+STAT3i-treated vs. vehicle-treated T-cell single-cell transcriptomes in PKT tumors. The horizontal bar in graded color from red (left) to blue (right) represents the enrichment of genes in the GSE41867 pathway, ranked from high expression in MEKi+STAT3i-treated (on the left) to high expression in vehicle-treated (on the right) datasets. PD-L1 levels were determined by **(B)** western blot of whole tumor lysate and **(C)** IHC analysis of vehicle, MEKi, STAT3i, or MEKi+STAT3i treated PKT mice. Data are shown as mean ± SEM. Scale bar = 50 μm.

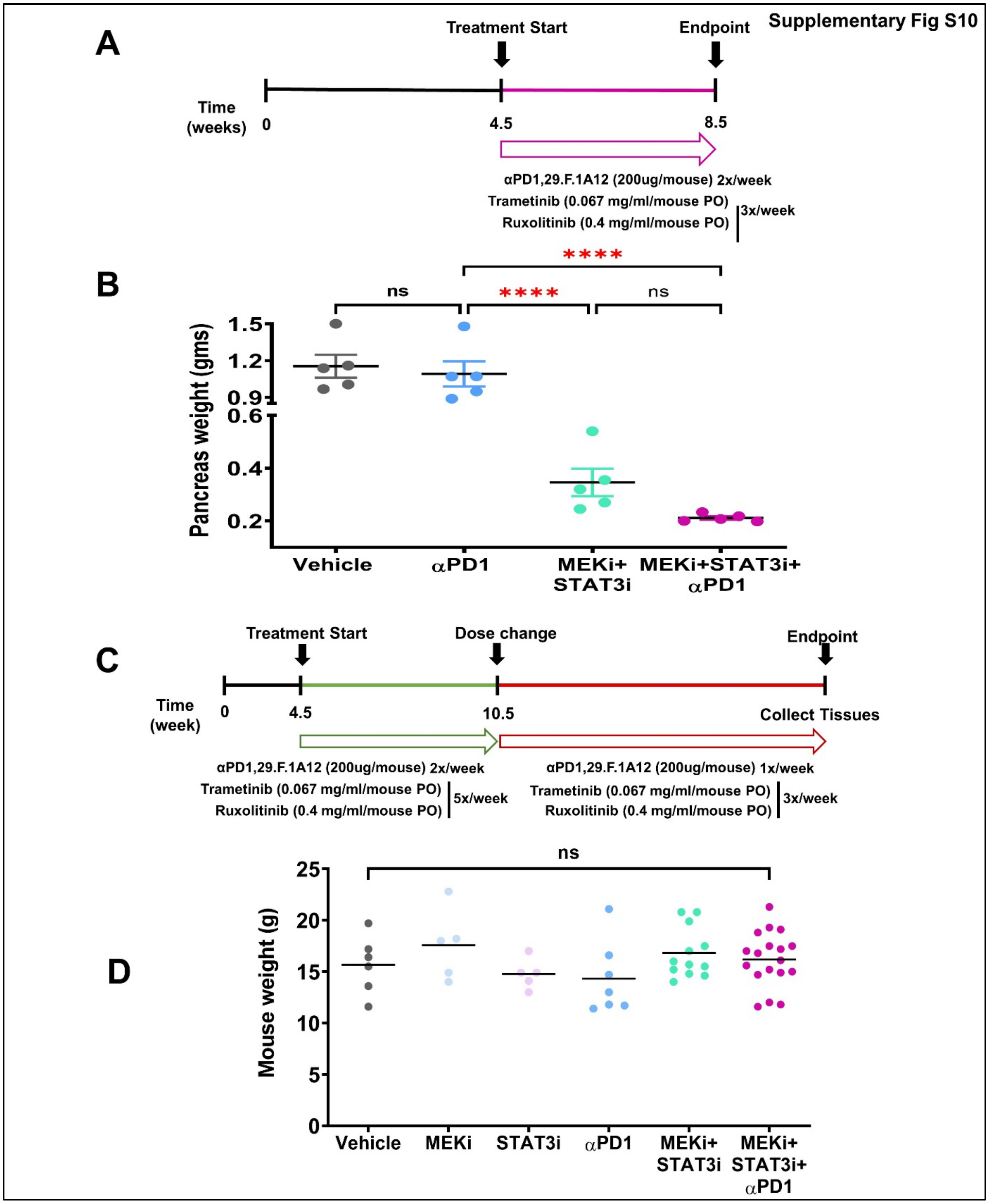


**Figure S10. Treatment schema for *in vivo* investigations in PKT mice for endpoint and survival experiments.** Treatment timeline and dosing schedule for **(A)** endpoint studies in PKT mice, and **(B)** pancreas weights in indicated treatment groups at designated endpoint. Treatment timeline and dosing schedule for **(C)** survival experiments in PKT mice, and **(D)** total mouse weights among treatment arms following 4 weeks of indicated therapies at endpoint analysis. Data are shown as mean ± SEM. ns, not significant, ****, P<0.0001.

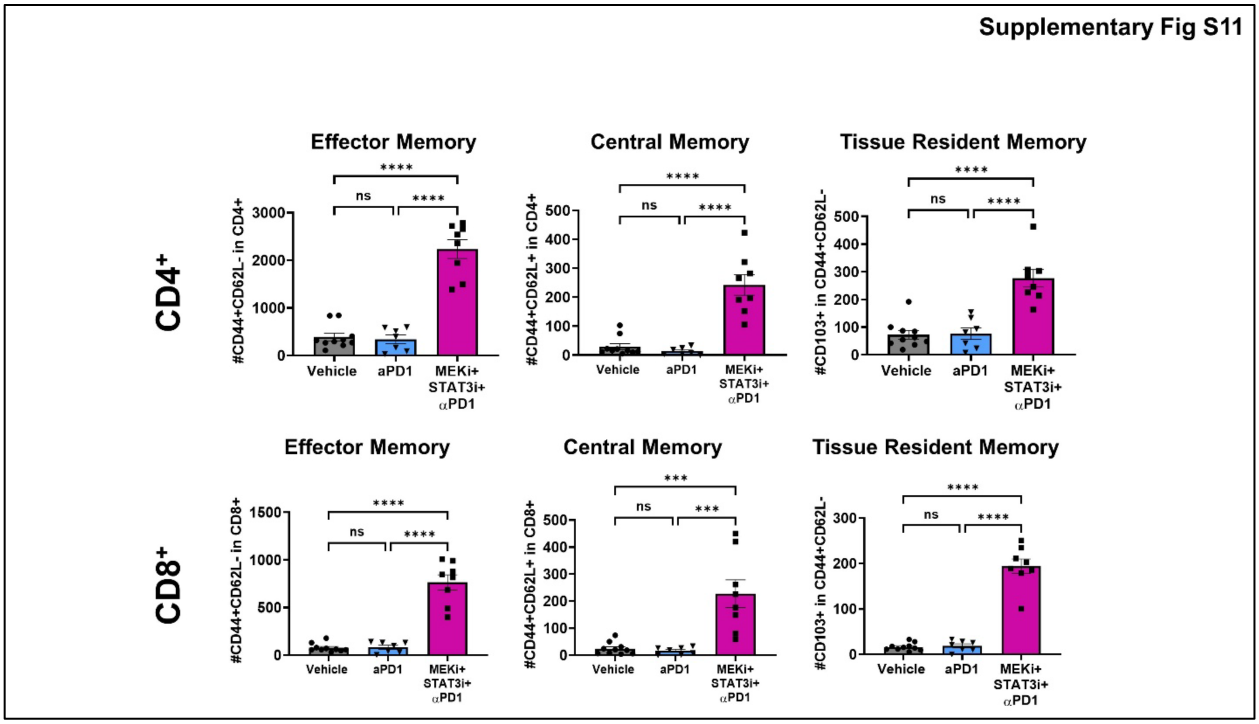


**Figure S11. Treatment with combined MEKi+STAT3i and αPD1 increases effector, central, and tissue-resident memory T-cell populations in the PDAC TME.** Levels of effector memory ($CD44^{+}CD62L^{-}$), central memory ($CD44^{+}CD62L^{+}$), and tissue-resident memory ($CD44^{+}CD62L^{-}CD103^{+}$) cells within $CD4^{+}$ (*top*) and $CD8^{+}$ (*bottom*) T-cell populations were determined by flow cytometry in PKT mice treated with vehicle, αPD1, and MEKi+STAT3i and αPD1 (n=7-8 mice/arm). Data are shown as mean ± SEM. ns, not significant; ***, p<0.001; ****, p<0.0001.

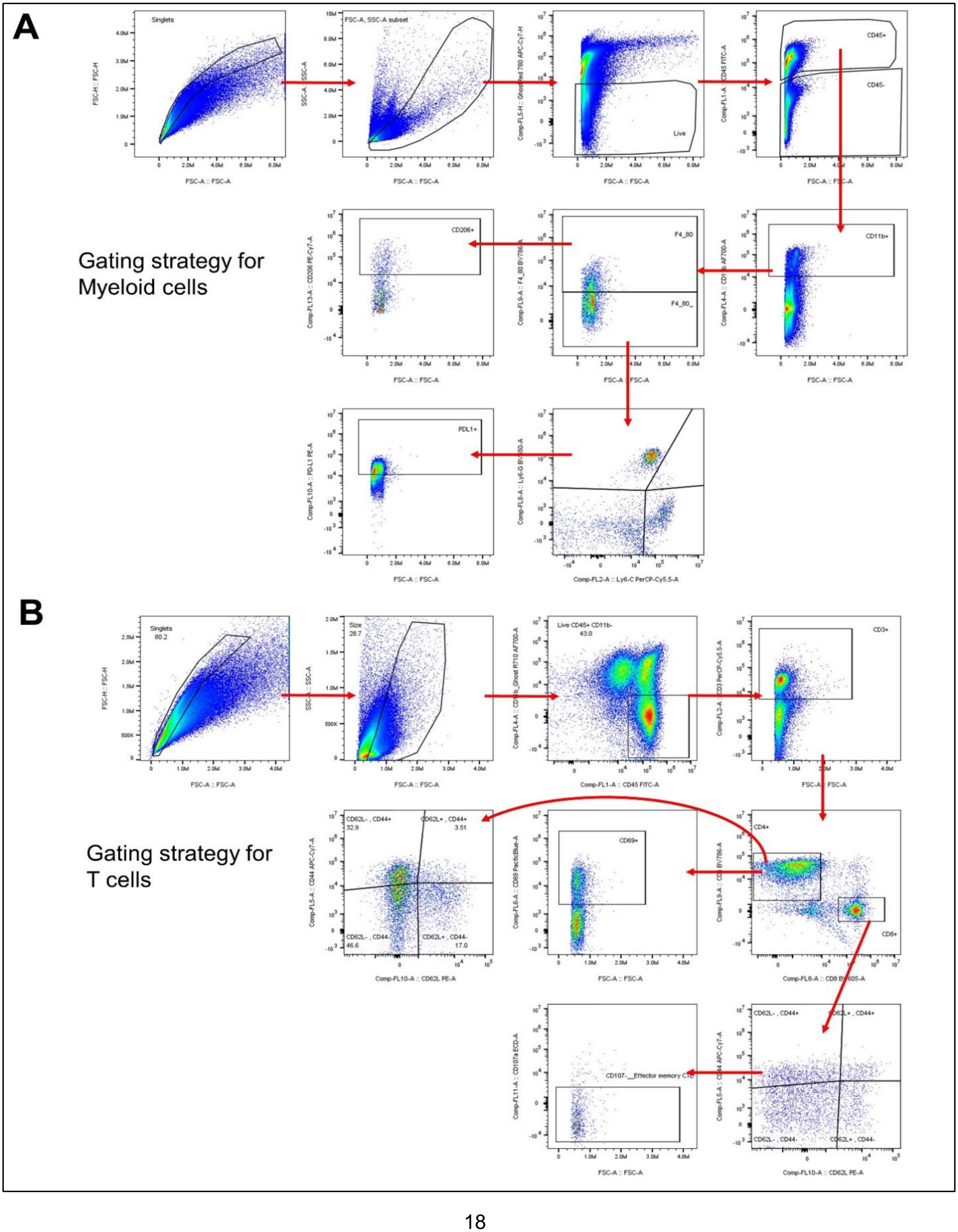
A
Gating strategy for Myeloid cells
Singlets
FSC-A, SSC-A subset
Live
CD45+
CD45-
CD11b+
F4_80
F4_80_
CD206+
PDL1+
B
Gating strategy for T cells
Singlets
80.2
Size
28.7
Live CD45+ CD11b-
43.0
CD3+
CD4+
CD8+
CD69+
CD62L- , CD44+
32.9
CD62L+ , CD44+
3.51
CD62L- , CD44-
46.6
CD62L+ , CD44-
17.0
CD107-__Effector memory CTL

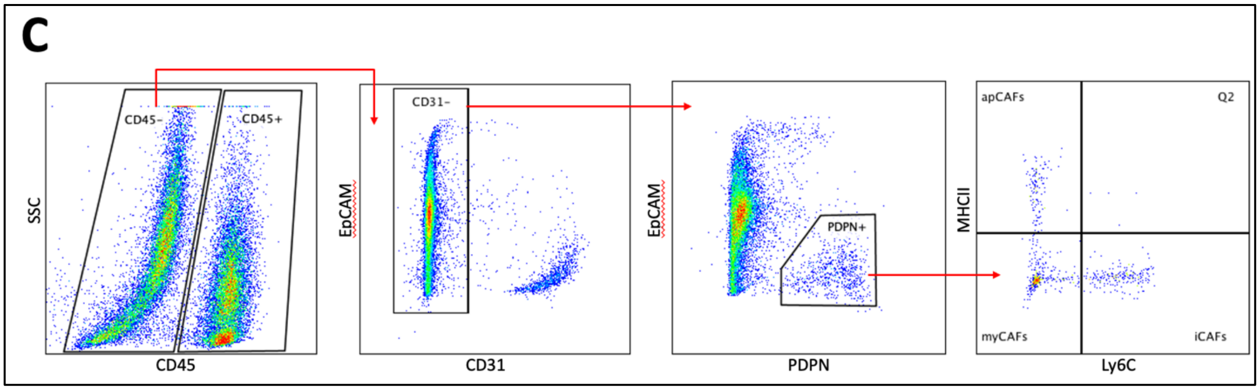


**Figure S12. Gating strategies used for flow cytometry analysis.** Representative diagrams depicting gating strategies for intratumoral **(A)** myeloid, **(B)** T-cell, and **(C)** CAF populations for flow cytometry experiments in PKT or orthotopic mice.

### SUPPLEMENTARY TABLES

**Table S1:** Primary antibodies for flow cytometry analysis

| Primary Antibodies | Fluorophore | Supplier | Species | Catalogue number |
|---|---|---|---|---|
| CD45 | FITC | Biolegend | Mouse | 103108 |
| CD3 | PerCPCy5.5 | Biolegend | Mouse | 100218 |
| CD4 | BV785 | Biolegend | Mouse | 100453 |
| CD8 | BV605 | Biolegend | Mouse | 100744 |
| CD44 | APC-Cy7 | Biolegend | Mouse | 103028 |
| CD62L | PE | Biolegend | Mouse | 104408 |
| CD69 | BV421 | Biolegend | Mouse | 104527 |
| PD-1 | PE-Cy7 | Biolegend | Mouse | 135216 |
| CD107a | PE-Dazzle | Biolegend | Mouse | 121624 |
| CD11b | AF700 | eBioscience | Mouse | 56-0112-82 |
| F4/80 | BV785 | Biolegend | Mouse | 123141 |
| Ly6G | BV650 | Biolegend | Mouse | 127641 |
| Ly6C | PerCPCy5.5 | Biolegend | Mouse | 128012 |
| CD206 | PE-Cy7 | Biolegend | Mouse | 141720 |
| Viability | Live/Dead Blue | Thermofisher | Mouse | L34962 |
| CD11b | BUV805 | BD Biosciences | Mouse | 741934 |
| PDPN | BV421 | Biolegend | Mouse | 127423 |
| CD45 | BV510 | Biolegend | Mouse | 103138 |
| MHCII | BV711 | Biolegend | Mouse | 107643 |
| EpCAM | PE | Biolegend | Mouse | 118206 |
| Ly6C | PE-Cy7 | Biolegend | Mouse | 128018 |
| CD31 | APC-Cy7 | Biolegend | Mouse | 102534 |

**Table S2:** Primary antibodies for CyTOF analysis

| Target | Supplier | Label | Clone | Reactivity |
|---|---|---|---|---|
| CD45 | Fluidigm | 89Y | 30-F11 | Mouse |
| Ly-6G/C (Gr-1) | Fluidigm | 141Pr | RB6-8C5 | Mouse |
| TCRb | Fluidigm | 143Nd | H57-597 | Mouse |
| CD69 | Fluidigm | 145Nd | H1.2F3 | Mouse |
| F4/80 | Fluidigm | 146Nd | BM8 | Mouse |
| CD44 | Fluidigm | 147Sm | IM7 | Mouse |
| CD19 | Fluidigm | 149Sm | 6D5 | Mouse |
| CD3e | Fluidigm | 152Sm | 145-2C11 | Mouse |
| CD8a | Fluidigm | 153Eu | 53-6.7 | Mouse |
| CD11b (Mac-1) | Fluidigm | 154Sm | M1/70 | Mouse |
| CD279 (PD-1) | Fluidigm | 159Tb | 29F.1A12 | Mouse |
| CD62L (L-selectin) | Fluidigm | 160Gd | MEL-14 | Mouse |
| CD278/ICOS | Fluidigm | 168Er | C398.4A | Mouse |
| CD206 (MMR) | Fluidigm | 169Tm | C068C2 | Mouse |
| CD80 (B7-1) | Fluidigm | 171Yb | 16-10A1 | Mouse |
| CD4 | Fluidigm | 174Yb | RM4-5 | Mouse |

**Table S3:** Guide RNA sequence information for CRISPR/Cas9 genetic editing of Mek1 and Stat3 in KPC CAFs

| Target Gene | Guide Sequence |
|---|---|
| **Mek1/Map2K1**<br>**Catalogue No:** CCMV1101-26395<br>**Vector-** pCLIP-gRNA-EFS-BlastV122 | sgRNAseq1# **AAGCACAAGATTATGCACAG**<br>sgRNAseq2# **GTACATCGTGGGCTTCTACG**<br>sgRNAseq3# **GTAGAAGCCCACGATGTACG**<br>Non-Targeting# **GGAGCGCACCATCTTCTTCA** |
| **Stat3**<br>**Catalogue No:** CCMV1101-20848<br>**Vector-** pCLIP-gRNA-EFS-PuroV71 | sgRNAseq1# **GAGATTATGAAACACCAACG**<br>sgRNAseq2# **CTGCTTCTCTGTCACTACGG**<br>sgRNAseq3# **GTTTACCACGAAAGTCAGGT**<br>Non-Targeting# **GGAGCGCACCATCTTCTTCA** |

**Table S4:** Primary antibodies for immunohistochemistry and immunofluorescence analysis

| **Primary Antibodies (IHC)** | **Supplier** | **Species** | **Catalogue number** |
|---|---|---|---|
| PD-L1 | Cell Signaling | Rabbit | 13684 |
| αSMA | Abcam | Rabbit | ab5694 |
| Ki67 | Cell Signaling | Rabbit | 9027 |

| **Primary Antibodies (IF)** | **Supplier** | **Species** | **Catalogue number** |
|---|---|---|---|
| CD11b | Abcam | Rat | ab8878 |
| F4/80 | Cell Signaling | Rabbit | 70076S |
| Ly6G | Abcam | Rat | ab25377 |
| CD31 | Cell Signaling | Rabbit | 77699S |
| Hoescht | Invitrogen | X | 33342 |

**Table S5:** Primary antibodies for Western Blot analysis

| **Primary Antibodies** | **Supplier** | **Species** | **Catalogue number** |
|---|---|---|---|
| PD-L1 | Cell Signaling | Rabbit | 13684 |
| pSTAT3 | Cell Signaling | Rabbit | 9145 |
| tSTAT3 | Cell Signaling | Mouse | 9139 |
| pERK | Cell Signaling | Rabbit | 4376 |
| tERK | Cell Signaling | Rabbit | 5013 |
| GAPDH | Cell Signaling | Rabbit | 2118 |
